# Apolipoprotein E O-glycosylation is associated with amyloid plaques and *APOE* genotype

**DOI:** 10.1101/2023.01.03.522616

**Authors:** Paige E Lawler, James G Bollinger, Suzanne E Schindler, Cynthia R Hodge, Nicolas J Iglesias, Vishal Krishnan, John B Coulton, Yan Li, David M Holtzman, Randall J Bateman

## Abstract

Although the *APOE* ε4 allele is the strongest genetic risk factor for sporadic Alzheimer’s disease (AD), the relationship between apolipoprotein (apoE) and AD pathophysiology is not yet fully understood. Relatively little is known about the apoE protein species, including post-translational modifications, that exist in the human periphery and CNS. To better understand these apoE species, we developed a LC-MS/MS assay that simultaneously quantifies both unmodified and O-glycosylated apoE peptides. The study cohort included 47 older individuals (age 75.6 ± 5.7 years [mean ± standard deviation]), including 23 individuals (49%) with cognitive impairment. Paired plasma and cerebrospinal fluid samples underwent analysis. We quantified O-glycosylation of two apoE protein residues – one in the hinge region and one in the C-terminal region – and found that glycosylation occupancy of the hinge region in the plasma was significantly correlated with plasma total apoE levels, *APOE* genotype and amyloid status as determined by CSF Aβ42/Aβ40. A model with plasma glycosylation occupancy, plasma total apoE concentration, and *APOE* genotype distinguished amyloid status with an AUROC of 0.89. These results suggest that plasma apoE glycosylation levels could be a marker of brain amyloidosis, and that apoE glycosylation may play a role in the pathophysiology of AD.

**Highlights:**

- Simultaneous quantification of unmodified and O-glycosylated apoE via LC-MS/MS.
- Total plasma apoE varies by *APOE* genotype in an isoform, dose-dependent fashion.
- CNS-derived apoE is more extensively O-glycosylated than peripherally derived apoE.
- Hinge region glycosylation occupancy of plasma apoE is associated with amyloidosis.

## 1. Introduction

As the most significant genetic risk factor for the development of late-onset Alzheimer’s disease (AD), apolipoprotein E (apoE protein, *APOE* gene) has been the focus of a considerable amount of biomedical research. The gene is polymorphic, with three common allelic isoforms derived from non-synonymous mutations in the codons for the amino acids at positions 112 and 158 [1]. The three isoforms, apoE2 (cys112, cys158), apoE3 (cys112, arg158), and apoE4 (arg112, arg158), vary subtly in amino acid sequence but substantially in terms of overall charge, structure, and function. While the *APOE* ε2 allele is associated with decreased risk for AD, the ε4 allele is associated with a dose-dependent, increased risk – 3-fold for heterozygotes and 12-fold for homozygotes – relative to the common *APOE* ε3 allele [2]–[7]. Despite this well-established association, a complete understanding of how the *APOE* alleles differentially modulate AD pathogenesis remains elusive. A significant challenge to this endeavor has been a lack of robust methods to fully identify and quantify the array of proteoforms that are found in the independently synthesized pools of apoE in the central nervous system and the periphery [8], [9].

The mature 299 amino acid apoE sequence folds to a tertiary structure composed of two distinct structural domains joined by a flexible hinge region (Figure 1). The N-terminal domain (amino acids 1-191) contains the receptor binding region, while the C-terminal domain (amino acids 225-299) contains the region responsible for binding lipids [7]. ApoE is expressed in the periphery by hepatocytes and, to a lesser extent, by macrophages and monocytes [7], [10]. It is expressed in the central nervous system (CNS) by microglia, astrocytes, vascular smooth muscle cells, the choroid plexus and, in a more limited context, by neurons under stress [11]—[16]. There is compelling evidence from liver transplant studies that apoE in the periphery and the CNS is generated from two independent metabolic pools [8]. Additionally, post-translational modification of apoE is differentially regulated in the periphery compared to the CNS [17]. All newly synthesized apoE has an 18-amino-acid signal peptide that is ultimately cleaved from the mature protein [18]. This peptide directs apoE to the Golgi apparatus and through the classical secretory pathway. During the secretory process, apoE is both glycosylated and sialylated [17]. The extent and amino acid location of these modifications is cell and tissue-specific and likely impacts the interaction of apoE with a variety of its receptors, ligands, and cofactors. Despite recent advances in both mass spectrometry and apoE structural elucidation, the impact and biological relevance of O-glycosylation remains unclear.

**Figure 1.**
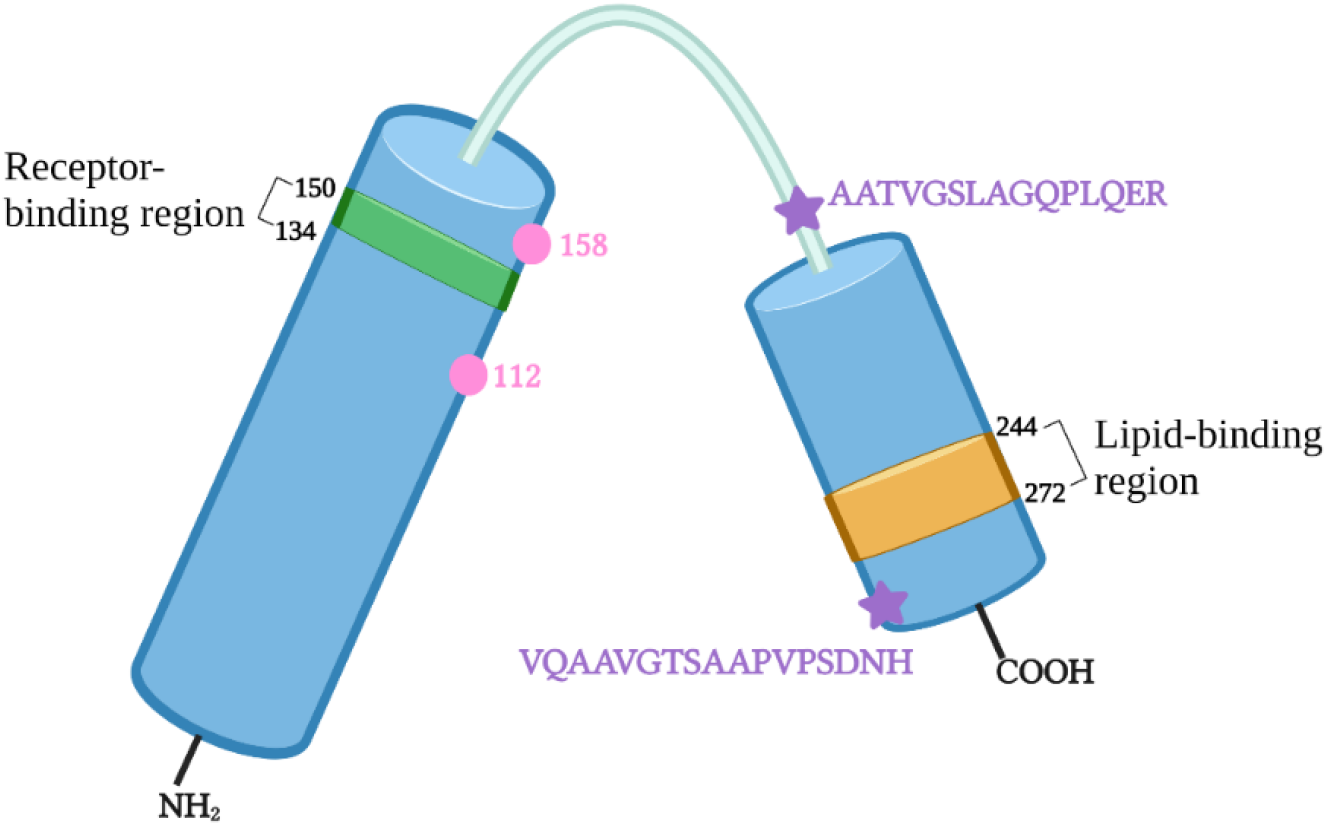
Illustration of the structural and functional regions of apolipoprotein E. The tertiary structure of apoE consists of two distinct structural domains joined by a hinge region. The receptor-binding region (green) is located in the N-terminal domain and spans residues 134 to 150. The lipid-binding region (orange) is located in the C-terminal domain and spans residues 244-272. The two isoform-specific residues of apoE are amino acids 112 and 158 (pink circles). The two O-glycosylated peptides characterized in this study are shown (purple stars).

This study describes the characterization and quantification of both unmodified and O-glycosylated apoE peptides in the plasma and cerebrospinal fluid samples of 47 older individuals (Table 1A,B). The goal of these experiments was two-fold. First, to characterize the complete apoE protein backbone and determine reported trends in absolute amounts of apoE, and second, to identify and quantify O-glycosylation, and compare glycosylation profiles across biofluids, *APOE* genotypes and amyloid statuses.

**Table 1.** Characteristics of samples studied sorted by amyloid status (A) and *APOE* genotype (B). Amyloid status was determined based on CSF amyloid beta 42/40 concentration ratio (amyloid positive if 42/40 ratio < 0.12). *APOE* ε4 carriers were defined by the presence of at least one ε4 allele (ε3/ε4, ε4/ε4) in contrast to *APOE* ε4 non-carriers (ε2/ε3, ε3/ε3). *APOE* genotype was confirmed by proteotyping via MS/MS. Clinical Dementia Rating (CDR) scores of 0, 0.5, and 1 correspond to cognitively unimpaired, very mild

| Characteristics | Amyloid –<br>(n = 23) | Amyloid +<br>(n = 24) | Entire cohort<br>(n = 47) |
| --- | --- | --- | --- |
| Age (years ± SD, range) | 74.6 ± 5.7<br>(65.9 – 87.7) | 77.3 ± 5.8<br>(63.6 – 85.8) | 75.9 ± 5.8<br>(63.6 – 87.7) |
| Sex (n, % Female) | 11, 48% | 11, 46% | 22, 47% |
| Race (n, %) |  |  |  |
| Black / African American | 1, 5% | 1, 4% | 2, 4% |
| White | 22, 95% | 23, 96% | 45, 96% |
| *APOE* ε4 carrier status (n) | Carrier = 5<br>Non-carrier = 18 | Carrier = 15<br>Non-carrier = 9 | Carrier = 20<br>Non-carrier = 27 |
| *APOE* genotypes (n) | ε2/ε3 = 3<br>ε3/ε3 = 15<br>ε3/ε4 = 5<br>ε4/ε4 = 0 | ε2/ε3 = 1<br>ε3/ε3 = 8<br>ε3/ε4 = 12<br>ε4/ε4 = 3 | ε2/ε3 = 4<br>ε3/ε3 = 23<br>ε3/ε4 = 17<br>ε4/ε4 = 3 |
| CDR (0/0.5/1 n, % >0) | 15/5/3, 35% | 9/12/3, 62% | 24/17/6, 49% |

**Table 1 Characteristics of samples studied sorted by amyloid status (A) and APOE genotype (B).**
| Characteristics | $\epsilon 2/\epsilon 3$<br>(n = 6) | $\epsilon 3/\epsilon 3$<br>(n = 21) | $\epsilon 3/\epsilon 4$<br>(n = 17) | $\epsilon 4/\epsilon 4$<br>(n = 3) |
| --- | --- | --- | --- | --- |
| Age (years ± SD, range) | 73.2 ± 8.9<br>(63.6 – 87.7) | 78.2 ± 4.5<br>(68.6 – 85.8) | 74.8 ± 5.6<br>(65.9 – 84.9) | 73.1 ± 6.2<br>(68.3 – 80.2) |
| Sex (n, % Female) | 2, 33% | 9, 43% | 10, 59% | 1, 33% |
| Race (n, %) |  |  |  |  |
| Black / African American | 1, 17% | 0, 0% | 1, 6% | 0, 0% |
| White | 5, 83% | 21, 100% | 16, 94% | 3, 100% |
| Amyloid status (n) | Amyloid – = 5<br>Amyloid + = 1 | Amyloid – = 13<br>Amyloid + = 8 | Amyloid – = 5<br>Amyloid + = 12 | Amyloid – = 0<br>Amyloid + = 3 |

## 2. Materials and Methods

### 2.1. Cohort Characteristics

Participants with plasma and cerebrospinal fluid samples collected at the same session were selected from the late-onset Alzheimer’s disease (LOAD)-100 cohort that has previously been described based on sample availability [19]. The study cohort included 47 older individuals (age 75.6 ± 5.7 years [mean ± standard deviation]), including 23 individuals (49%) with cognitive impairment as defined by a Clinical Dementia Rating (CDR) of 0.5 or greater. Amyloid status was determined based on CSF amyloid beta 42/40 (amyloid positive if 42/40 < 0.12): 23 individuals were amyloid negative and 24 were amyloid positive. In the entire cohort, 47% of participants were female, 96% identified as White, and 43% carried at least one *APOE* ε4 allele.

### 2.2. Production of a biomimetic analytical standard

ApoE3* and apoE4* isotopically labelled internal standards were obtained from the media of immortalized astrocytes derived from transgenic mice expressing human *APOE* ε3 or *APOE* ε4 [20]. Briefly, separate astrocyte cultures were grown in L-[U-^13^C_6_,^15^N_4_] arginine-enriched, serum-free media to produce secreted lipoparticles containing heavy arginine-labeled apoE3* and apoE4*. These media samples were combined and cross-titrated with unlabeled media internal standards to establish equivalent concentrations between apoE3* and apoE4*. The equimolar concentration of apoE3* and apoE4* in the standard allows for the correction of any potential isoform-specific biases that could be introduced in sample preparation or analysis.

### 2.3. Immunoprecipitation and tryptic digestion of apoE

ApoE was immunoprecipitated from matched participant samples of 20 μL CSF and 2 μL plasma via a monoclonal antibody to human apoE (HJ15.3) (provided by the Holtzman Lab) conjugated to M-270 Epoxy Dynabeads (Life Technologies/Invitrogen, Carlsbad, CA). The CSF samples were prepared by combining 20 μL of neat CSF with 100 μL of the apoE3*/apoE4* astrocyte media internal standard, 100 μL of a ^15^N-labeled recombinant apoE3 internal standard (Promise Proteomics, Grenoble, France) and 750 μL of a PBS solution containing a cocktail of protease inhibitors. The plasma samples were prepared by first making a 1:100 sample dilution (10 μL plasma, 990 μL PBS) and then combining 200 μL of said dilution with 100 μL of apoE3*/apoE4* media standard, 100 μL of recombinant apoE internal standard and 570 μL of PBS/PI buffer solution. 30 μL of the HJ15.3 antibody-bead slurry was added to each sample and the samples were immunoprecipitated on a KingFisher Flex instrument. Briefly, following a one hour mixing step, the beads were washed three times (once with 1X PBS and twice with 50 mM TEABC) and the target was eluted off the beads in 100 μL neat formic acid. The eluent was dried using positive pressure heated nitrogen, treated with 100 μL acetonitrile, and then dried again. Resultant samples were stored at −20°C overnight. Then, each sample was heat denatured and reduced via a 0.5-hour, 37°C incubation in 20 μL of a 0.1 M DTT/40% TFE solution, and alkylated via a 0.5-hour, room temperature incubation in 0.618 μL of 20 mM iodoacetamide. The alkylation was quenched with 10 μL 0.1 mM DTT. Samples were diluted with 170 μL of 100 mM TEABC and then digested at 37°C for 4 hours with 1 μG trypsin. Prior to LC-MS/MS analysis, solid-phase extraction was performed on the samples using the Oasis mixed-mode cation exchange sorbent following the manufacturer’s instructions.

### 2.4. LC-MS/MS analysis

Following isolation, digestion, and solid-phase extraction clean-up, each digest was subjected to nano-flow liquid chromatography and analysis via tandem mass spectrometry in which a previously developed targeted proteomic assay monitoring 19 unmodified apoE tryptic peptides and 10 glycopeptides was applied. Analyses were performed on a Thermo TSQ Altis Triple Quadrupole mass spectrometer (Thermo Fisher Scientific, Waltham MA, USA) interfaced with a Waters M-Class nanoAcquity chromatography system (Waters, Milford, MA. USA). Extracted digests were reconstituted in 25 μL of 0.1% formic acid in water. Reconstituted digests were loaded via direct injection (4.5 μL) onto a Waters 100 x 0.075 mm nanoEase M/Z HSS C18 T3 capillary column (Waters, Milford, MA. US) at 0.5% acetonitrile/0.1% formic acid in water with a flow rate of 600 nL/min for 12 minutes. After loading, peptides were resolved using a 15-minute linear gradient at 300 nL/min from 0.5% acetonitrile/0.1% formic acid in water to 30% acetonitrile/0.1% formic acid in water. The initial gradient was followed by a steeper linear gradient to 60% acetonitrile/0.1% formic acid over 2 minutes at 300 nL/min. The column was then held at this solvent composition for an additional 2 minutes at the same flow rate. Following a 7-minute wash of the column with 95% acetonitrile/0.1% formic acid in water at 600 nL/min, the column was equilibrated back to initial solvent conditions for 5 minutes at 600 nL/min. Peaks were detected using Xcalibur and quantified, reviewed and edited using Skyline (MacCoss Lab, University of Washington Genome Sciences Department). All pertinent data collection parameters as well as a list of peptide precursor and product ions used for each monitored SRM transition is provided in Supplementary Table 1.

### 2.5. Absolute quantification of unmodified apoE peptides

Absolute amounts of each apoE peptide are reported as a ratio of the ^14^N peak area to the corresponding peak area of the ^15^N-labeled recombinant apoE internal standard for the given peptide. The results are plotted as a function of peptide amino acid start position and each point represents the mean for the given group, with error bars indicating the SEM. To compare absolute apoE amounts overall, the reported ^14^N/^15^N ratios for all common peptides were summed and an average calculated for each sample.

### 2.6. Relative quantification of unmodified apoE peptides

Relative amounts of each apoE peptide are calculated by taking the “absolute” amount as a ratio to the “absolute” amount of the reference peptide LGPLVEQGR. The results are plotted as a function of peptide amino acid start position and each point represents the mean for the given group, with error bars indicating the SEM. This approach allowed for the localization of peptides where the native protein may differ across different groups or biofluids. Relative calculations were also done with the peptide GEVQAMLGQSTEELR and correlation plots between the two were constructed to increase confidence that the reference peptide used did not introduce bias (Supplementary Figure 8).

### 2.7. O-glycosylation site occupancy calculation

Six unique glycoforms of the peptide AATVGSLAGQPLQER were detected and quantified. Five unique glycoforms of the peptide VQAAVGTSAAPVPSDNH were detected and quantified. The ^14^N peak areas for unmodified and all O-glycosylated species of the given peptide (AATVGSLAGQPLQER or VQAAVGTSAAPVPSDNH) were summed and the site occupancy of each glycoform was calculated as a ratio of the ^14^N peak area of the glycoform to the summed ^14^N peak area. Total site occupancy was calculated by summing the site occupancies of each glycoform.

### 2.8. Statistical analysis

Data was visualized using GraphPad Prism software (GraphPad, San Diego, CA, USA, v9.4.0). Unpaired T-tests and one-way ANOVA (followed by Tukey post hoc T-tests when applicable) were performed for group comparisons. Results were further evaluated using analysis of covariance (ANCOVA) models in which age was treated as a continuous covariate and amyloid status, *APOE* ε4 carrier status or *APOE* genotype, and sex were treated as classification predictors. These analyses were implemented in PROC GLM/SAS. Receiver operating characteristic (ROC) analyses were performed to evaluate the ability of plasma glycosylation occupancy combined with plasma total apoE amount and *APOE* genotype to predict amyloid status and were implemented with PROC LOGISTIC.

## 3. Results

### 3.1. Absolute quantification of apoE peptide profile

To fully characterize the apoE peptide backbone, apoE was measured across 19 common peptides (excluding isoform-specific peptides), covering 67% of the protein amino acid sequence, in plasma and CSF. In the plasma, comparisons across *APOE* genotypes revealed an isoform-dependent, dose-dependent trend in the total amount of apoE (ε2/ε3 > ε3/ε3 > ε3/ε4 > ε4/ε4). A one-way Analysis of Variance (ANOVA) demonstrated a statistically significant difference across groups (p < 0.0001), with post hoc pairwise comparisons indicating significant differences between the ε2/ε3 and ε3/ε3, ε2/ε3 and ε3/ε4, and ε2/ε3 and ε4/ε4 groups (Figure 2A,C). This isoform-dependent trend remained after adjusting for amyloid status, age and sex (p = 0.0004), (Supplementary Table 1), as well as when the comparison was made by *APOE* ε4 carrier status (p = 0.002) (Supplementary Figure 1A). Comparing absolute amounts of apoE across *APOE* genotypes in the CSF, there was not a clear isoform- or dose-dependent trend. Instead, ε3/ε3 and ε3/ε4 demonstrated a similar protein profile and trended towards higher absolute amounts of apoE compared to the ε2/ε3 and ε4/ε4 groups, but the differences were not statistically significant by one-way ANOVA (p = 0.139) (Figure 2B,D), even after adjusting for variables including amyloid status, age and sex (Supplementary Table 2). Comparisons across *APOE* ε4 carrier status were also not significant in the CSF (p = 0.646) (Supplementary Figure 1B). Total plasma and CSF apoE levels did not vary by amyloid status (Supplementary Figures 2A-D) (p = 0.239 for plasma; p = 0.608 for CSF). Additionally, plasma and CSF apoE amounts were not correlated with one another, including after adjusting for variables including amyloid status, age and sex (Supplementary Table 3).

**Figure 2.**
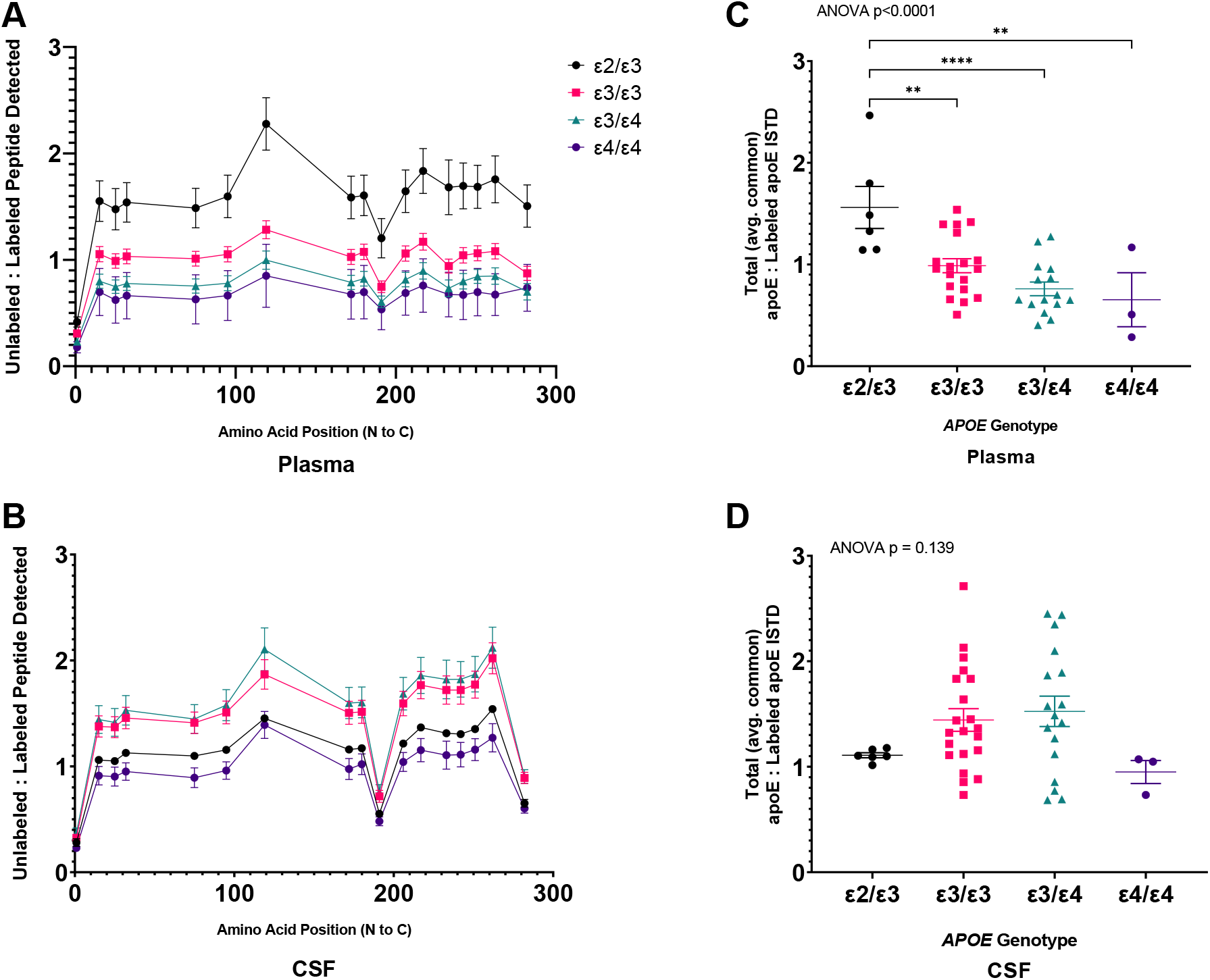
Absolute quantification of the apoE peptide backbone indicates that amounts of plasma apoE decrease in an isoform-, dose-dependent fashion as ε2/ε3 > ε3/ε3 > ε3/ε4 > ε4/ε4, while CSF apoE amounts do not appear to follow a clear isoform- or dose-dependent trend. **A,B** Each point represents the mean value for the corresponding peptide (plotted along the X-axis by amino acid start position) for all samples of a given *APOE* genotype in the plasma (**A**) and CSF (**B**), with error bars indicating the SEM. **C,D** Each point represents the average of all peptide values for a given participant in the plasma (**C**) and CSF (**D**), with error bars indicating the SEM. Asterisks designate statistically significant differences between *APOE* genotype groups.

### 3.2. Relative quantification of apoE peptide profile

Comparing the relative levels of apoE peptides between plasma and CSF, there were two peptides that differed significantly: AATVGSLAGQPLQER (amino acid positions 192-206; p < 0.0001) and VQAAVGTSAAPVPSDNH (amino acid positions 283-299; p < 0.0001) (Figure 3). For both peptides, the relative amount was lower in the CSF than in the plasma. Comparisons across *APOE* genotypes (Supplementary Figure 3A,B) and across amyloid status (Supplementary Figure 4A,B) did not reveal any significant differences in the relative levels of any of the peptides in the plasma or in the CSF.

**Figure 3.**
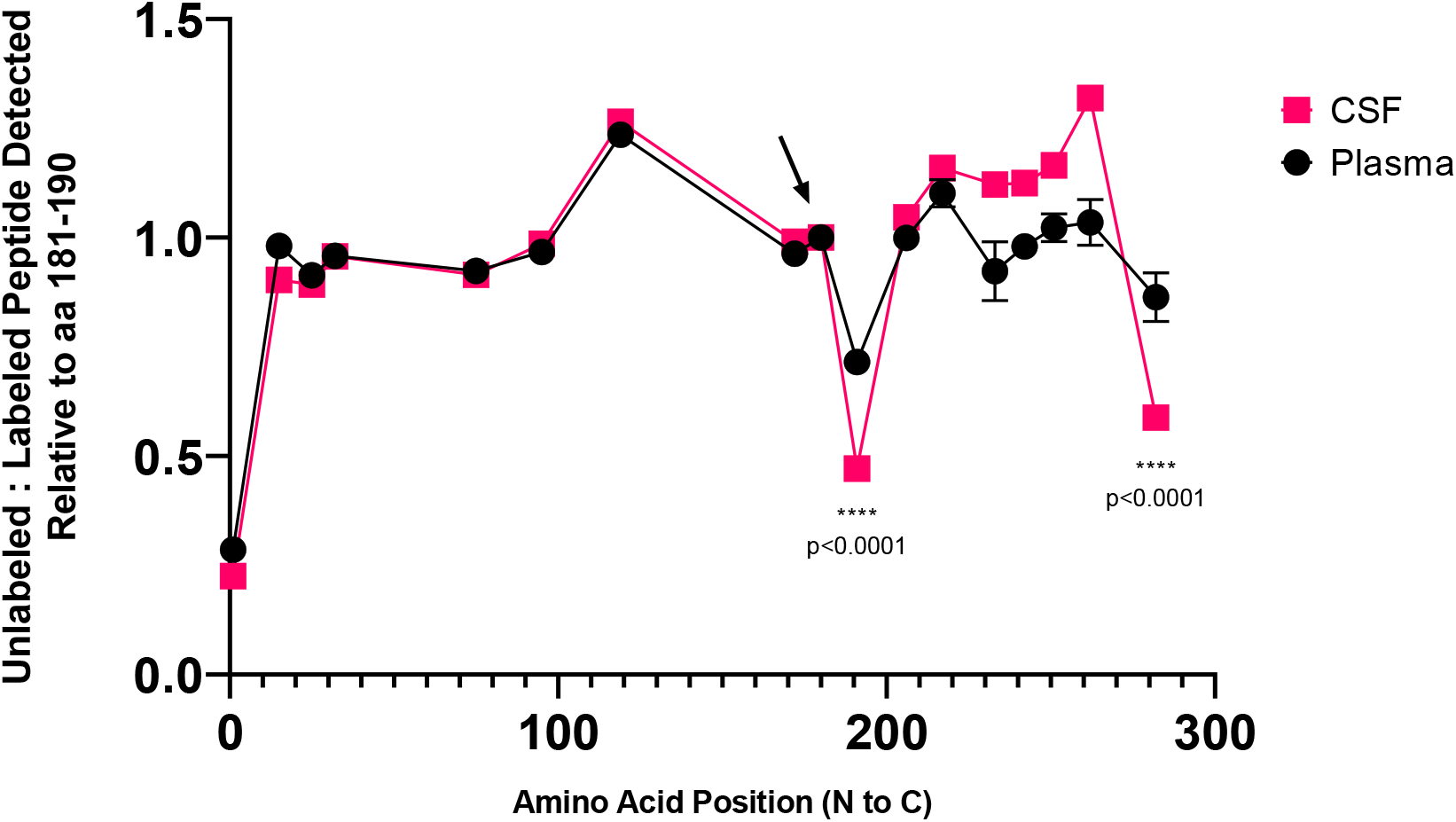
Comparison of the relative amounts of apoE across the unmodified backbone reveal differences between the plasma and the CSF at positions 192-206 and 283-299. Each point represents the mean value for the corresponding peptide relative to the reference peptide LGPLVEQGR (denoted with a black arrow) for all plasma (black circle) and CSF (pink square) samples, with error bars indicating the SEM. Asterisks designate the peptides that show a statistically significant difference between the plasma and CSF samples, where the p-value was less than 0.0001 as determined by a paired sample T-test.

### 3.3. Characterization of apoE O-glycopeptides

For the qualitative assessment of apoE O-glycosylation, we leveraged access to biomimetic stable isotope-labeled analytical standards and high-resolution tandem mass spectrometry. The biomimetic standards were used in addition to the uniformly ^15^N-labeled recombinant apoE3 standard to enhance confidence in the identification of specific glycopeptides (Supplementary Figures 5 and 6). Additionally, diagnostic oxonium ions derived from collision induced fragmentation of these glycopeptides were used to further increase confidence in their characterization [21].

### 3.4. Characterization and quantification of apoE O-glycopeptides in the hinge region

The quantitative assessment of O-glycosylation of the hinge region peptide, AATVGSLAGQPLQER, revealed a significant difference in the total O-glycosylation site occupancy between the plasma and CSF, with the plasma apoE carrying 4.3% modification compared to 10.6% modification of CSF apoE (p < 0.0001) (Figure 4A). Additionally, the distribution of glycospecies differed between the biofluids (Figure 4B). Of the three most abundant glycospecies in the plasma, the predominant structure was the monosialylated core 1 (2.1%), followed by the disialylated core 1 (1.2%) and the unsialylated core 1 (0.07%). In contrast in the CSF, the predominant structure was the disialylated core 1 (5.7%), followed by the monosialylated core 1 (2.9%) and the unsialylated core 1 (1.1%). Total glycosylation site occupancy of the hinge region peptide was evaluated separately in the plasma and the CSF as a function of total apoE amount and across *APOE* genotype, amyloid status and cognitive status (Figure 5, Figure 6A-F). In both the plasma and the CSF, there was an overall moderate negative correlation between total site occupancy and total apoE amount, though the correlation did not reach significance in the CSF (*ρ* = −0.32 and p = 0.035 in plasma; *ρ* = −0.22 and p = 0.13 in CSF) (Figure 5). Of all comparisons across *APOE* genotype, amyloid status and cognitive status in the plasma and the CSF, the only one which reached significance was the comparison by *APOE* genotype in the plasma (p = 0.044) Figure 6A-F, Figure 7). An Analysis of Covariance (ANCOVA) demonstrated that in the plasma, total site occupancy was negatively associated with plasma total apoE amount (p = 0.0006), lower in *APOE* ε4 carriers (p < 0.0001), and higher in amyloid positive individuals (p = 0.003), even after adjusting for age and sex (Table 2A,B, Supplementary Figure 7). Additional ANCOVAs performed using the site occupancy of each of the three predominant glycoforms as the outcome variables revealed that the unsialylated core 1 and monosialylated core 1 species showed the strongest association with plasma total apoE amount, *APOE* ε4 carrier status and amyloid status (Supplementary Table 4). Receiver operating characteristic analysis of a model with plasma total site occupancy, plasma total apoE amount, and *APOE* genotype gave an area under the curve (AUC) of 0.89 (95% confidence intervals [CI] 0.80-0.98) for amyloid status (Figure 8). In contrast, the same analysis in the CSF showed no significant association of total site occupancy with total CSF apoE amount, or by *APOE* ε4 carrier status or amyloid status. Notably, there did appear to be an association of total O-glycosylation site occupancy with sex (p = 0.006) and age (p = 0.003) (Supplementary Table 5A,B).

**Figure 4.**
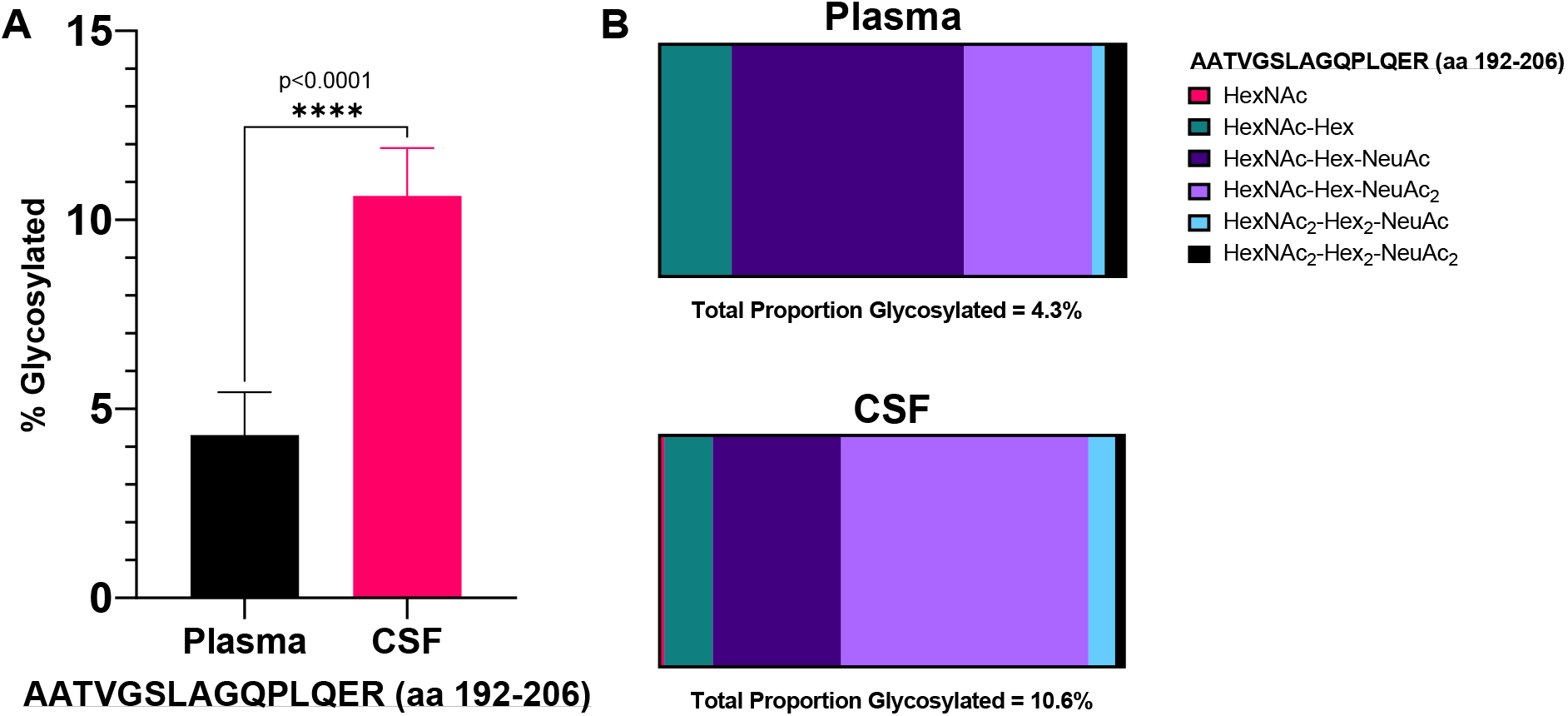
Comparison of plasma and CSF O-glycosylation site occupancy of the hinge region peptide, AATVGSLAGQPLQER, reveals that the CSF (10.6%) has more than twice the modification of plasma (4.3%), and a different distribution of modifications, including more relative and absolute amounts of the disialylated core 1 structure. **A** Total site occupancy in the plasma versus the CSF graphed as a percent of the total peptide amount. Asterisks designate a statistically significant difference between the groups, where the p-value was less than 0.0001 as determined by a paired sample T-test. **B** Relative distribution of glycospecies in the plasma and CSF, with colors indicating relative amounts of O-glycosylation structures quantified.

**Figure 5.**
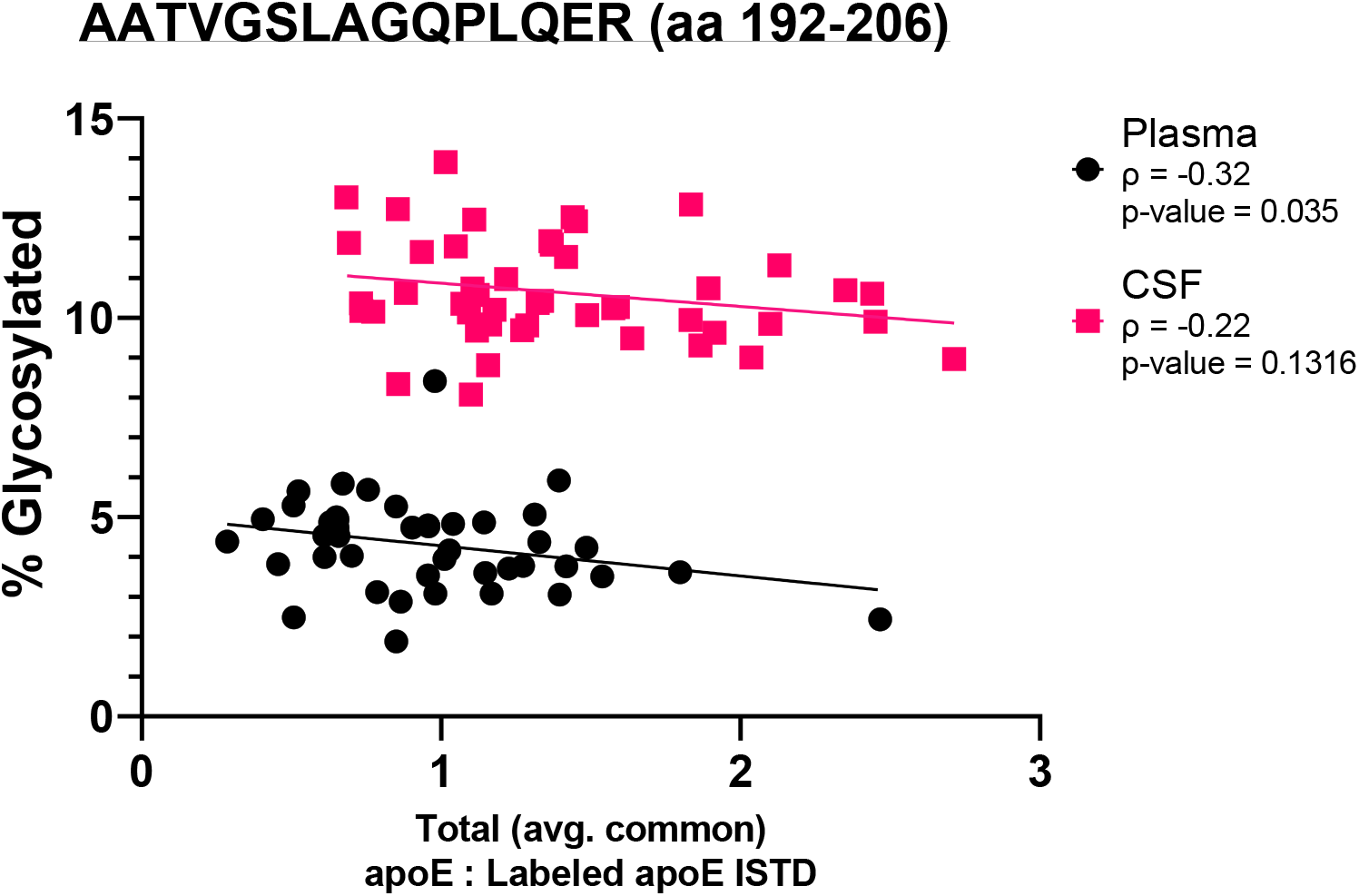
Percent O-glycosylation site occupancy of the hinge region peptide, AATVGSLAGQPLQER, as a function of total apoE amounts in the plasma and CSF. Each point represents the percent O-glycosylation site occupancy plotted against the corresponding total apoE amount (calculated as the average of all peptide values) for each sample. Spearman’s correlation coefficients and p-values are indicated.

**Figure 6.**
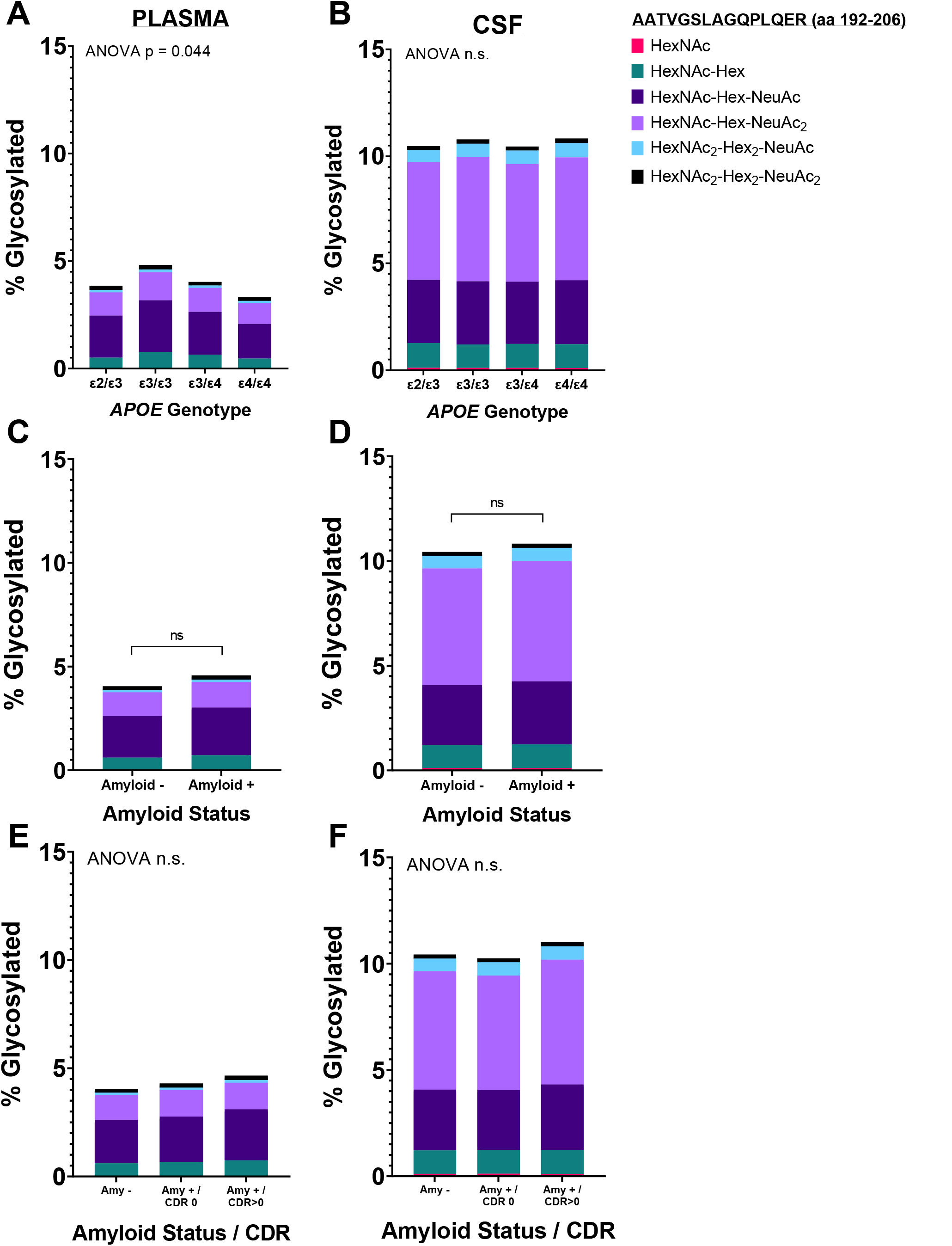
The total site occupancy and relative distribution of O-glycosylation modifications of the hinge region peptide, AATVGSLAGQPLQER, were compared between *APOE* genotypes, the presence of amyloid plaques in the brain and clinical dementia status in plasma and CSF. Differences in total O-glycosylation site occupancy by *APOE* genotype in the plasma were significant (p = 0.044). Total site occupancy was not significantly different by amyloid or cognitive status in the plasma, or by any comparisons in the CSF. The relative distribution of modifications was not significantly different across any comparisons in the plasma or CSF. The left panel shows results from the plasma and the right panel, the CSF. **A, B** Results compared across *APOE* genotypes. **C, D** Results compared across amyloid status. **E, F** Results compared across amyloid and cognitive status (determined by CDR score). Statistical significance was measured one-way ANOVA and unpaired T-tests where appropriate.

**Figure 7.**
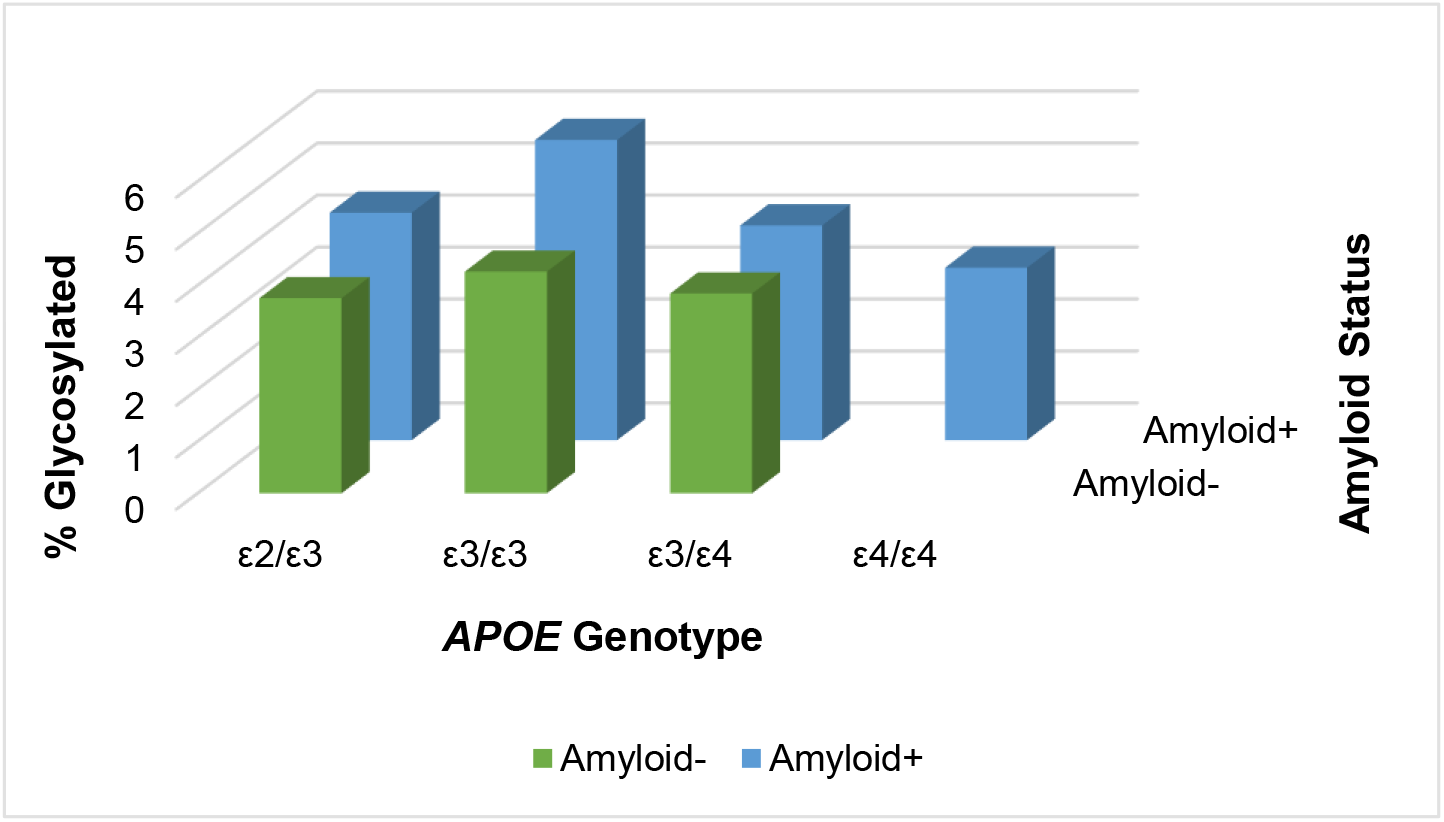
Plasma total O-glycosylation site occupancy of the hinge region peptide is correlated with APOE genotype and amyloid status.

**Figure 8.**
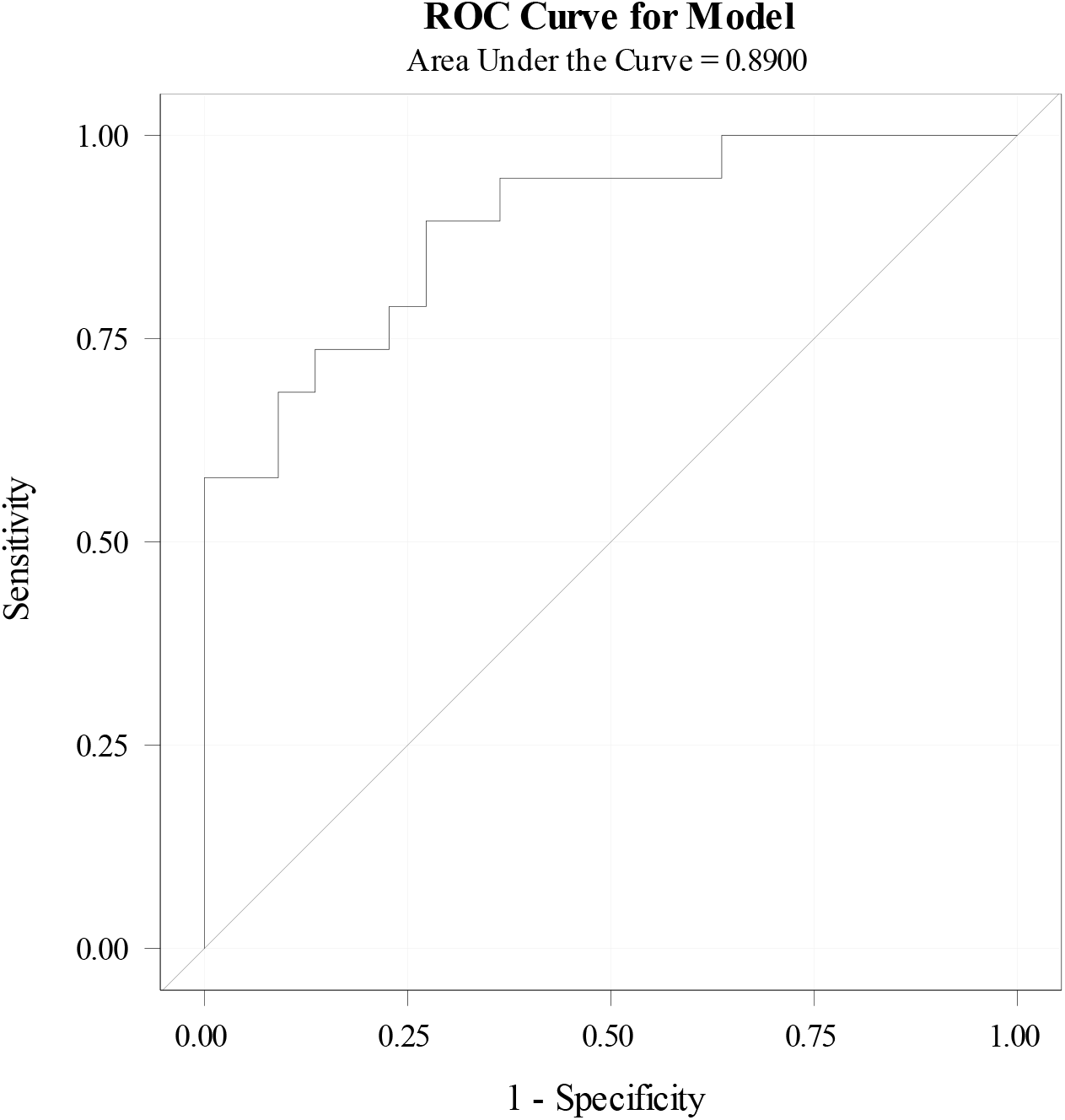
Receiver operating characteristic analysis demonstrates that a model including plasma O-glycosylation site occupancy of the hinge region peptide, plasma total apoE and *APOE* genotype may be predictive of amyloid status (AUC = 0.89). The area under the curve is noted with 95% confidence intervals (0.80-0.98).

**Table 2.** ANCOVA of plasma total O-glycosylation site occupancy of the hinge region peptide reveals a significant correlation with plasma total apoE amount (p = 0.0006), *APOE* ε4 carrier status (p < 0.0001), and amyloid status (p = 0.003). **A** Total site occupancy as the outcome variable with plasma total apoE as a continuous covariate and *APOE* ε4 carrier status and amyloid status as categorical covariates. **B** Total site occupancy as the outcome variable with plasma total apoE as a continuous covariate and *APOE* ε4 carrier status, amyloid status, sex and age as covariates.

| Parameter | Estimate | SE | *p* = |
| --- | --- | --- | --- |
| Intercept | 0.058 | 0.005 | <.0001 |
| Plasma_apoE | -0.014 | 0.004 | 0.0006 |
| *APOE* ε4 + | -0.015 | 0.003 | <.0001 |
| *APOE* ε4 - | 0 | . | . |
| Amyloid + | 0.009 | 0.003 | 0.003 |
| Amyloid - | 0 | . | . |

**Table 2 ANCOVA of plasma total O-glycosylation site occupancy of the hinge region peptide reveals a significant correlation with plasma total apoE amount (*p* = 0.0006), *APOE* ε4 carrier status (*p* < 0.0001), and amyloid status (*p* = 0.003).**
| Parameter | Estimate | SE | *p* = |
| --- | --- | --- | --- |
| Intercept | 0.036 | 0.021 | 0.084 |
| Plasma_apoE | -0.013 | 0.004 | 0.001 |
| *APOE* ε4 + | -0.014 | 0.004 | 0.0006 |
| *APOE* ε4 - | 0 | . | . |
| Amyloid + | 0.008 | 0.003 | 0.014 |
| Amyloid - | 0 | . | . |
| SEX-M | -0.0002 | 0.003 | 0.93 |
| SEX-F | 0 | . | . |
| AGE | 0.0003 | 0.0003 | 0.29 |

### 3.5. Characterization and quantification of apoE O-glycopeptides in the C-terminal region

The quantitative assessment of O-glycosylation of the C-terminal region peptide, VQAAVGTSAAPVPSDNH, revealed a significant difference in the total glycosylation site occupancy between the plasma and CSF, with the CSF carrying four-times (22.0%) the modification of the plasma (5.4%) (p < 0.0001) (Figure 9A). Additionally, as in the hinge region, the distribution of glycospecies differed between the biofluids (Figure 9B). In the plasma, the predominant structure was the monosialylated core 1 (3.9%), followed by the disialylated core 1 (1.0%) and the unsialylated core 1 (0.2%). In the CSF, the predominant structure was the disialylated core 1 (12.0%), followed by the monosialylated core 1 (7.5%) and the unsialylated core 1 (1.5%). Total glycosylation site occupancy of the C-terminal region peptide was evaluated separately in the plasma and the CSF as a function of total apoE amount and across *APOE* genotype, amyloid status and cognitive status (Figure 10, Figure 11A-F). In both the plasma and the CSF, there was an overall moderate positive correlation between percent glycosylation and total apoE amount (*ρ* = 0.17, p = 0.28 in plasma; *ρ* = 0.41, p = 0.0038 in CSF) (Figure 10), though the correlation did not reach significance in the plasma. Comparisons across *APOE* genotype, amyloid status and cognitive status did not reach significance for any analyses in either the plasma or CSF (Figure 11A-F). This remained largely the case in ANCOVA analyses (Supplementary Tables 6,7), although there was significant association of plasma total O-glycosylation site occupancy of the C-terminal region peptide with *APOE* ε4 carrier status (p = 0.021).

**Figure 9.**
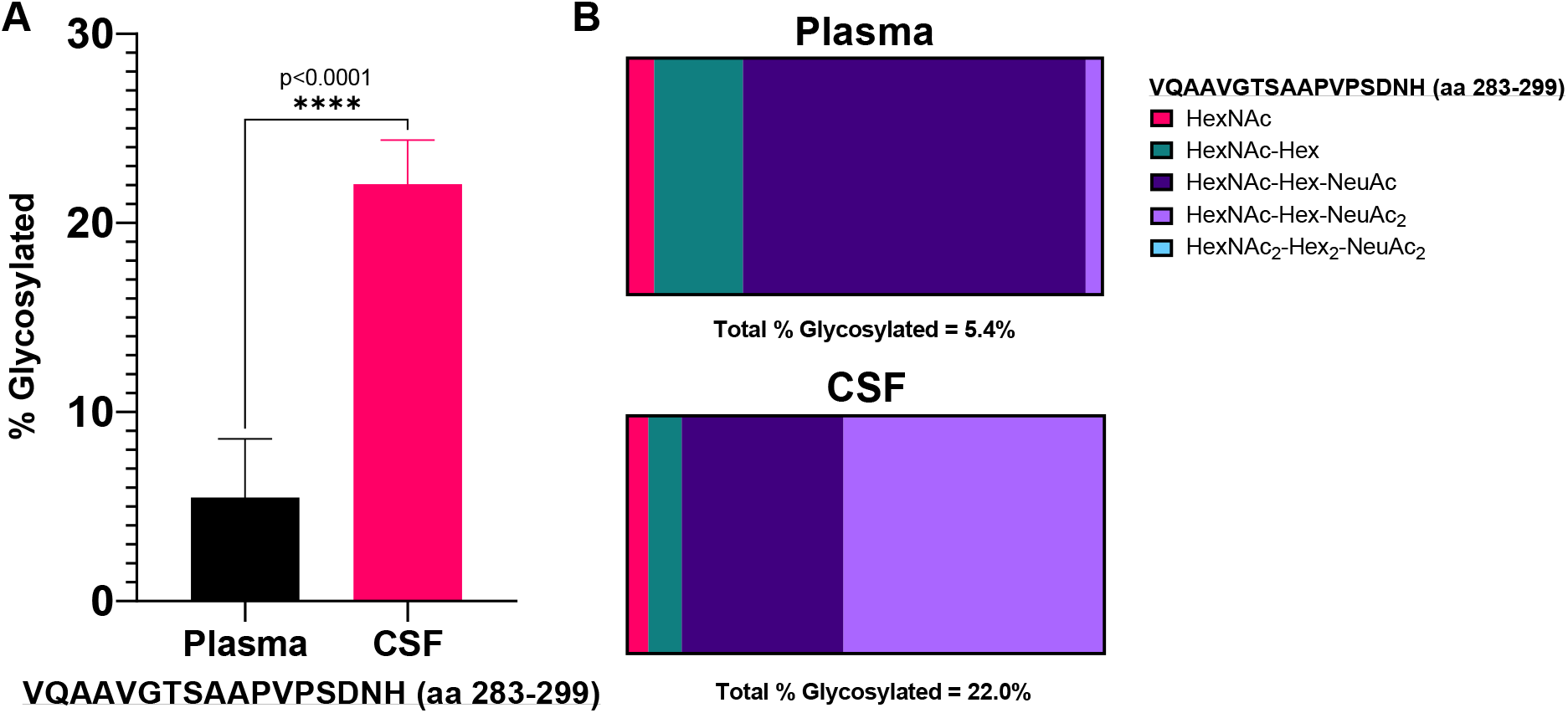
Comparison of plasma and CSF O-glycosylation site occupancy of the C-terminal peptide, VQAAVGTSAAPVPSDNH, reveals that the CSF (22.0%) has four-times the modification of plasma (5.4%), and a different distribution of modifications, including more relative and absolute amounts of the disialylated core 1 structure. **A** Total site occupancy in the plasma versus the CSF graphed as a percent of the total peptide amount. Asterisks designate a statistically significant difference between the groups, where the p-value was less than 0.0001 as determined by a paired sample T-test. **B** Relative distribution of glycospecies in the plasma and CSF, with colors indicating relative amounts of O-glycosylation structures quantified.

**Figure 10.**
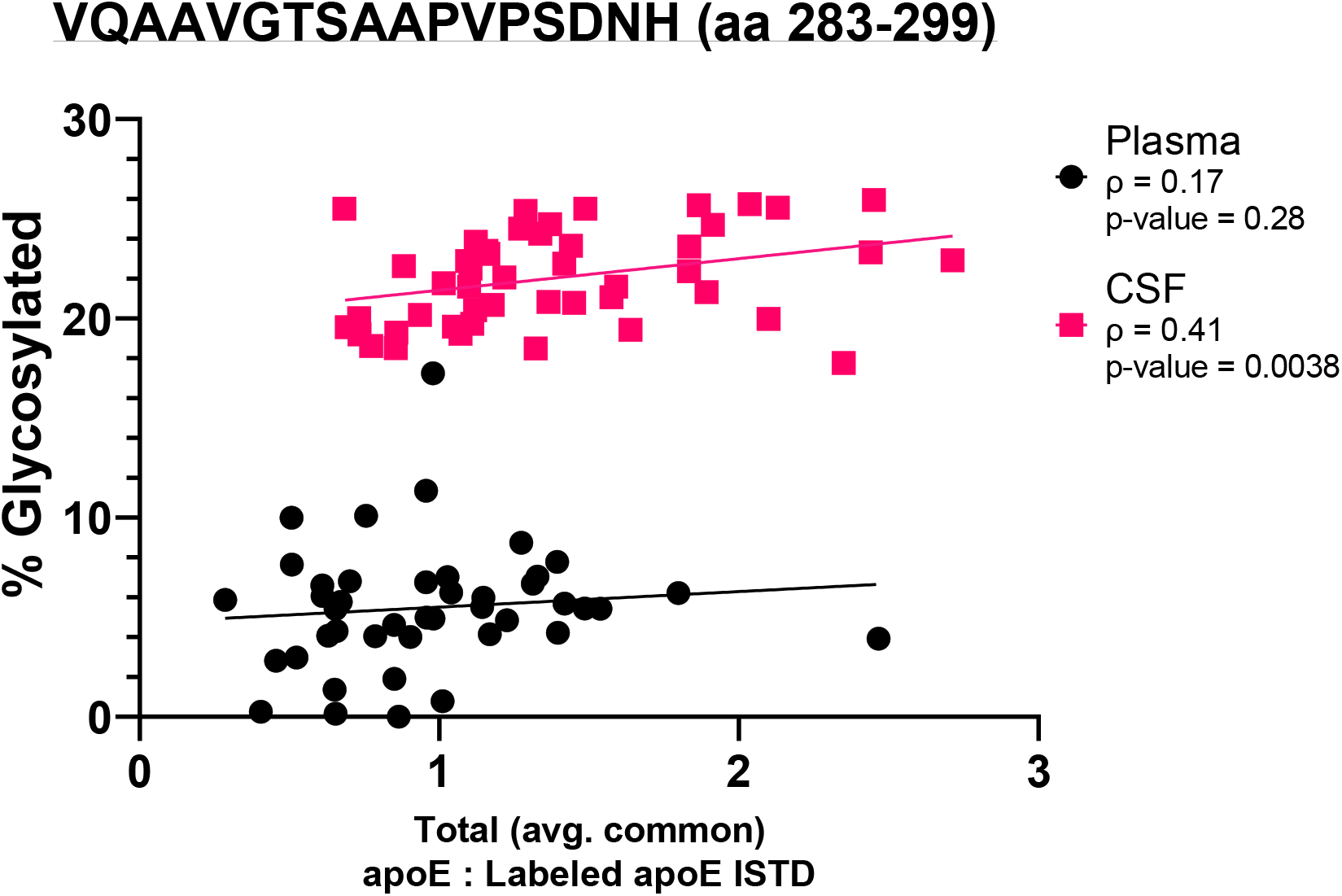
Percent O-glycosylation site occupancy of the C-terminal region peptide, VQAAVGTSAAPVPSDNH, is linearly positively correlated with total apoE amounts in the plasma and in the CSF. Each point represents the percent O-glycosylation site occupancy plotted against the corresponding total apoE amount (calculated as the average of all peptide values) for each sample. Rho Spearman’s correlation coefficients and p-values are indicated.

**Figure 11.**
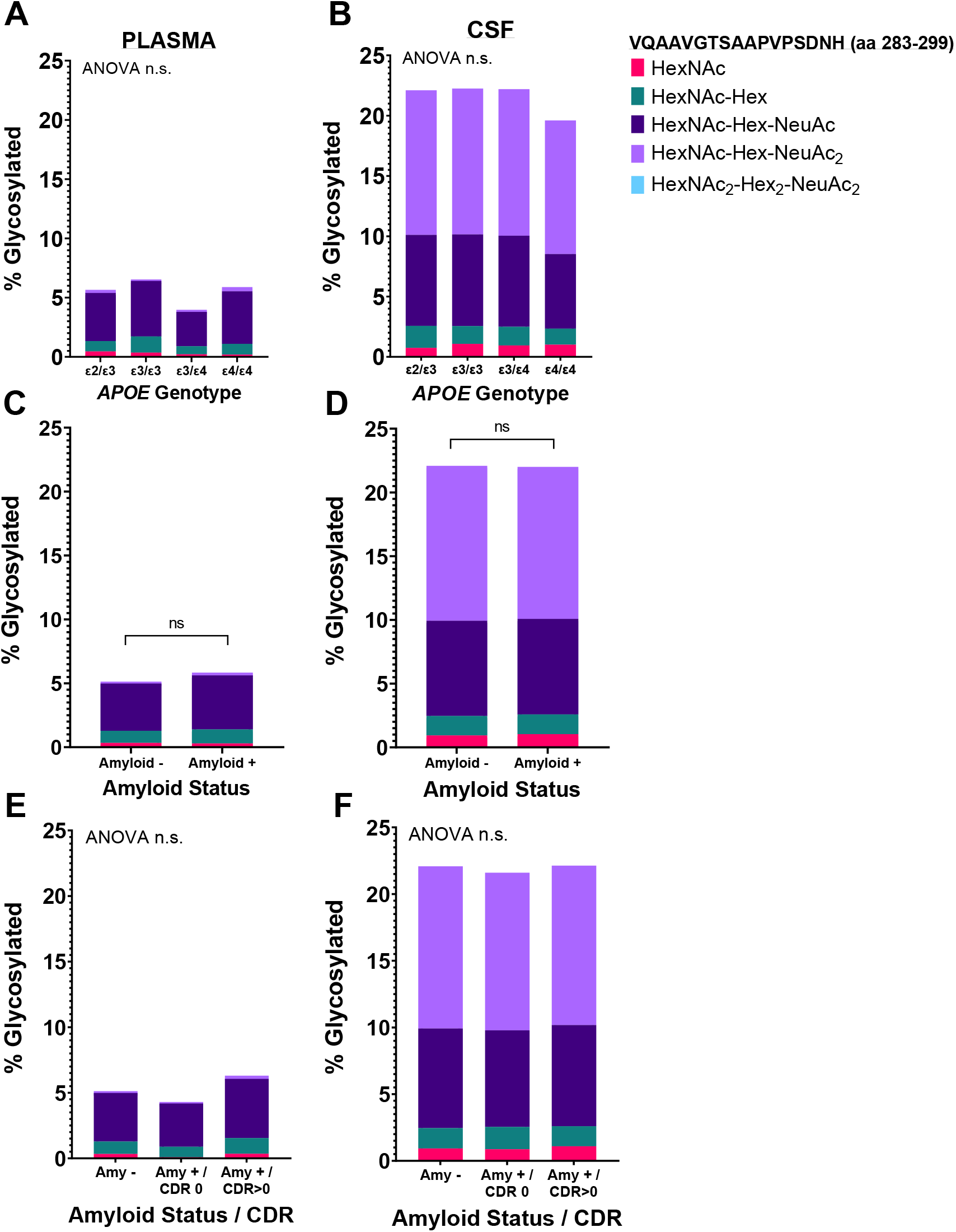
The total site occupancy and relative distribution of O-glycosylation modifications of the C-terminal peptide, VQAAVGTSAAPVPSDNH, were compared across *APOE* genotypes, the presence of amyloid plaques in the brain and clinical dementia status in plasma and CSF. Total site occupancy was not significantly different by any comparisons in the plasma or CSF. The relative distribution of modifications was not significantly different across any comparisons in the plasma or CSF. The left panel shows results from the plasma and the right panel, the CSF. **A, B** Results compared across *APOE* genotypes. **C, D** Results compared across amyloid status. **E, F** Results compared across amyloid and cognitive status (determined by CDR score). Statistical significance was measured one-way ANOVA and unpaired T-tests where appropriate.

## 4. Discussion

The results of this analysis confirm prior reports that apoE concentration in the periphery varies in an isoform, dose-dependent manner, and provide additional evidence that centrally derived apoE does not follow an isoform or dose-dependent trend in concentration. While the trend in the plasma is one that has been repeatedly reported in the literature [20], [22], [23], human studies of CSF apoE levels have yielded more conflicting results. Several groups have reported that apoE concentration in the CSF does vary in an isoform-dependent manner, notably that *APOE* ε2 is associated with higher concentrations while *APOE* ε4 is associated with lower concentrations: ε2 > ε3 > ε4 [24]–[27], while others report that CSF apoE levels do not vary according to *APOE* genotype [28], [29]. The inconsistencies in these findings could potentially be explained by differences in cohort design or assay accuracy and precision. For example, in prior immunoassays used to quantify apoE, such as ELISA, there could be an antibody bias for the different apoE isoforms. Conversely, the tandem mass spectrometry method used for the analyses in this study provides a more robust, non-antibody dependent quantitation method.

Seminal work on apoE glycosylation identified Thr194 as a major carbohydrate attachment site, identified glycosylations with variable amounts of sialic acid end-capping, and confirmed that glycosylation is not required for the secretion of apoE into the extracellular environment [30]. Subsequent efforts identified residues Thr8, Thr18, Thr189, Ser290, and Ser296 as additional attachment sites and suggested mucin type core 1 O-glycans as the predominant type of carbohydrate on apoE [31]–[34]. Our study did not detect the N-terminal glycosylations, which we attribute to the low amounts of plasma (2 μL) and CSF (20 μL) used. For both the hinge region and C-terminal glycosylations (Thr189, Ser290 and Ser296), glycopeptides were characterized using a combination of DIA and PRM analyses on a high-resolution tandem mass spectrometer. The distinction between the monosialylated and disialylated core 1 glycosylated peptides is apparent by their differences in retention time, precursor mass, and the relative intensity of the peaks at *m/z* 274.09 and 292.10 associated with sialic acid and the *m/z* 366.14 and 204.09 associated with the HexNAc-Hex and HexNAc saccharides, respectively. Additionally, the lability of the sialic acid bond leads to a high amount of in-source fragmentation of sialylated glycoforms [35], [36]. We addressed this by ensuring that our di-, mono-, and unsialylated glycopeptides are chromatographically resolved from one another [36]–[39].

Our analysis of O-glycosylation in the hinge and C-terminal regions of the apoE protein provides a detailed characterization and highly precise quantification of the amounts and types of modifications present in the plasma and CSF and across *APOE* genotypes, amyloid status and cognitive status. This study confirms that apoE is extensively glycosylated and sialylated, and that the extent of these modifications differs substantially in peripherally versus centrally derived pools of apoE, with higher absolute and relative CSF apoE modifications in both the hinge and C-terminal regions [31], [33]. The finding that CSF apoE holds substantially more glycosylation and sialylation than plasma apoE is particularly interesting in the context of lipoprotein particle binding, given previous studies that have demonstrated that the additional negative charge of sialylated glycans may contribute to improved HDL-binding [40], [41].

Perhaps the most salient finding from these analyses was that after adjusting for plasma total apoE, plasma glycosylation occupancy of the hinge region peptide showed a strong negative association with *APOE* ε4 carrier status (p < 0.0001) and a strong positive correlation with amyloid status (p = 0.003) (Table 2). These findings suggest a potentially mechanistic role of apoE between genotype status and amyloid deposition.

The finding that plasma glycoform levels vary significantly by genotype and amyloid status suggests that glycosylation occupancy of plasma apoE could potentially be a marker of brain amyloidosis and may have utility if incorporated into a blood-based biomarker assay for AD diagnosis. For example, a model including the plasma hinge region glycoform levels, plasma total apoE amount, and *APOE* genotype, distinguished amyloid status with an AUC of 0.89 (Figure 8). This model outperforms current *APOE* models, such as those incorporating *APOE* ε4 carrier status and age [42]. Given the relatively small size of the cohort studied, particularly the lower numbers of samples in the *APOE* ε2/ε3 and *APOE* ε4/ε4 groups, future studies to replicate this finding in other cohorts are needed.

Taken together, the results of this study provide further evidence that the independent pools of peripherally and centrally derived apoE appear to undergo distinct modification processes. Given the undeniable importance of *APOE* to AD, and the still limited understanding of how *APOE* affects risk for AD, studies of apoE metabolism are important. In addition to follow-up investigations of the hinge region glycosylation status of plasma apoE in other cohorts, future investigations may include analysis of brain tissue and further characterization of the glycoforms reported here. These findings will help to inform on the structural, and potentially functional, relationships of apoE to AD pathophysiology.

## CRediT authorship contribution statement

**Paige E Lawler:** Conceptualization; Data curation; Formal analysis; Roles/Writing - original draft. **James G Bollinger, PhD:** Conceptualization; Data curation; Formal analysis; Funding acquisition; Project administration; Writing - review & editing. **Suzanne E Schindler, MD, PhD:** Data curation; Formal analysis; Writing - review & editing. **Cynthia R Hodge, MBA:** Project administration; Writing - review & editing; Resources. **Nicolas J Iglesias:** Conceptualization; Writing - review & editing. **Vishal Krishnan, MD:** Conceptualization; Writing - review & editing. **John B Coulton, PhD:** Formal analysis; Writing – review & editing. **Yan Li, PhD:** Data curation; Writing - review & editing. **David M Holtzman, MD:** Conceptualization; Investigation; Methodology; Writing - review & editing. **Randall J Bateman, MD:** Conceptualization; Data curation; Formal analysis; Funding acquisition; Investigation; Methodology; Project administration; Resources; Supervision; Writing - review & editing

## Acknowledgements

We would like to express our gratitude to the research volunteers who participated in the studies from which these data were obtained and their supportive families. We would like to thank Hong Jiang and the Holtzman Lab for preparing the monoclonal antibody used for apoE analysis. We would also like to thank all members of the Bateman Lab for their continued support.

## Data availability

Data are available to qualified investigators who have an approved proposal.

## Appendix A. Supplementary information

**Supplementary Figure 1.**
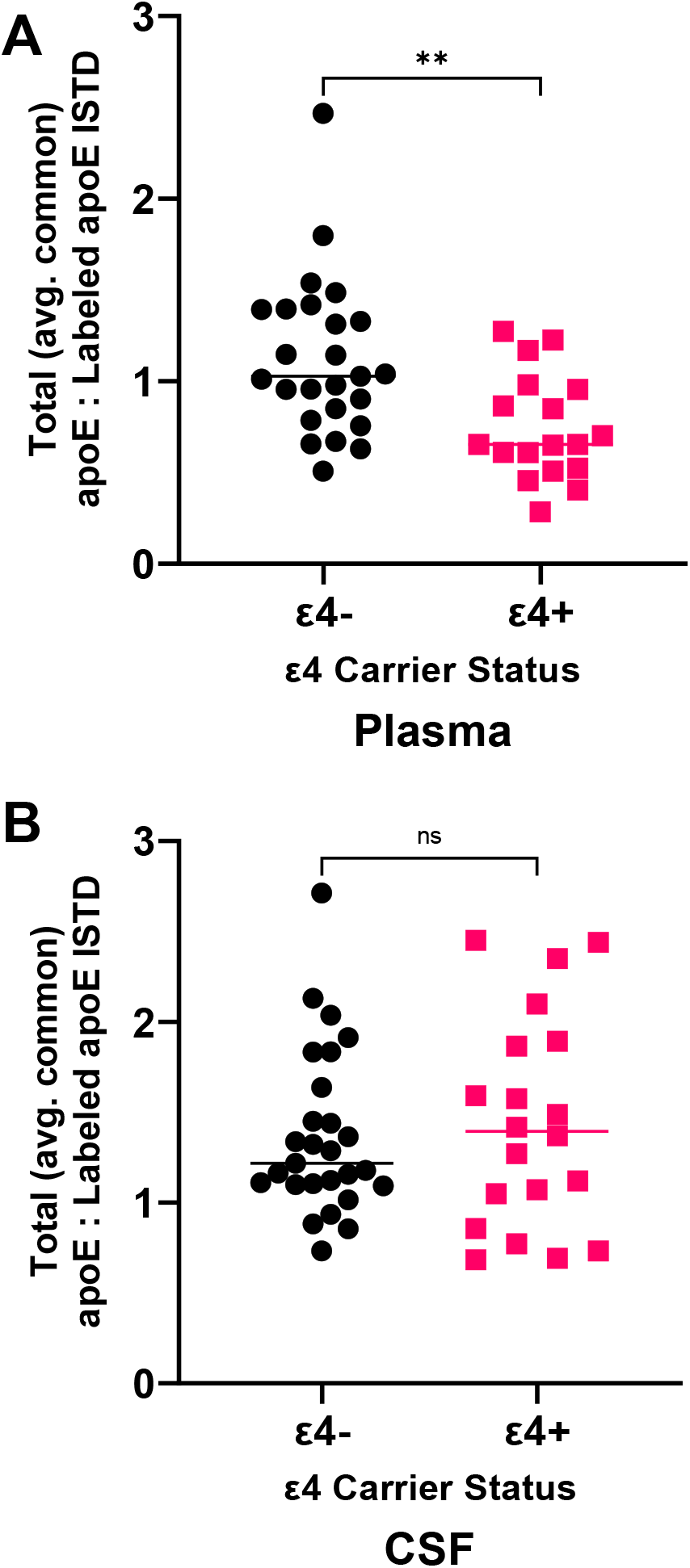
Absolute quantification of apoE indicates that amounts of plasma apoE vary by *APOE* ε4 carrier status, with ε4 carriers having lower total amounts of apoE compared to non-carriers, while amounts of CSF apoE do not differ significantly by ε4 carrier status. Each point represents the average of all peptide values for a given participant in the plasma (**A**) and CSF (**B**). Asterisks designate statistically significant differences between groups as determined by unpaired T-tests.

**Supplementary Table 1.** ANCOVA of plasma total apoE amount reveals a significant correlation with APOE genotype but not amyloid status, age or sex. The overall significance of *APOE* genotype was p = 0.0004.

| Source | Type III SS | *p* = |
| --- | --- | --- |
| *APOE* | 2.90 | 0.0004 |
| Amyloid | 0.043 | 0.55 |
| AGE | 0.0007 | 0.94 |
| SEX | 0.107 | 0.35 |

**Supplementary Table 2.** ANCOVA of CSF total apoE amount reveals no correlation with *APOE* genotype, amyloid status, age or sex. The overall significance of *APOE* genotype was p = 0.30.

| Source | Type III SS | *p* = |
| --- | --- | --- |
| *APOE* | 0.88 | 0.30 |
| Amyloid | 0.00007 | 0.98 |
| AGE | 0.40 | 0.20 |
| SEX | 0.035 | 0.70 |

**Supplementary Table 3.** ANCOVA reveals no correlation between CSF total apoE amount and plasma total apoE, after adjusting for *APOE* genotype, amyloid status, age and sex. The overall significance of *APOE* genotype was p = 0.22.

| Source | Type III SS | *p* = |
| --- | --- | --- |
| Plasma_apoE | 0.36 | 0.22 |
| *APOE* | 1.13 | 0.20 |
| Amyloid | 0.008 | 0.85 |
| AGE | 0.24 | 0.31 |
| SEX | 0.13 | 0.46 |

**Supplementary Figure 2.**
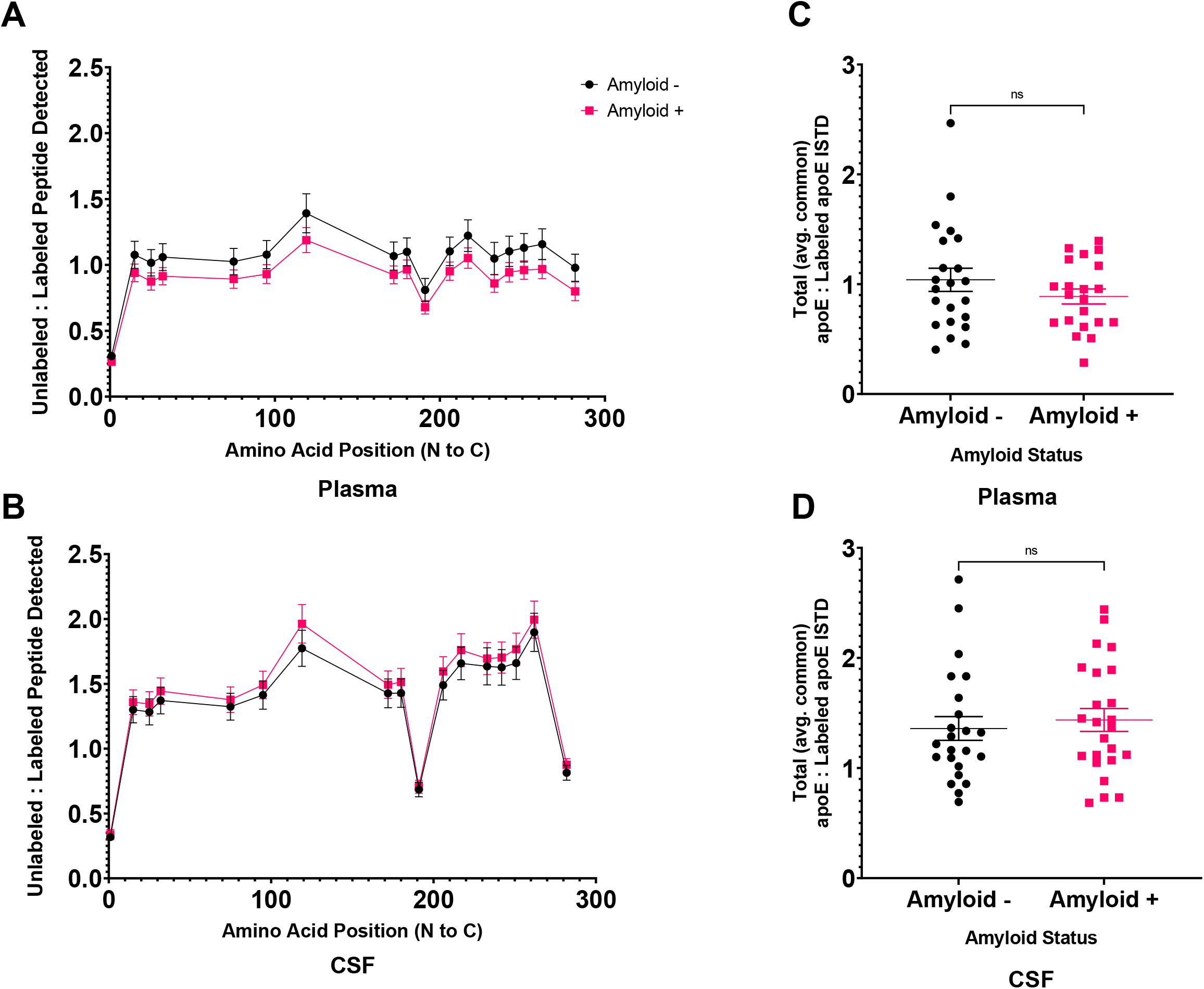
Absolute quantification of the apoE peptide backbone indicates that amounts of apoE are not significantly different across amyloid status in the plasma or in the CSF. **A,B** Each point represents the mean value for the corresponding peptide (plotted along the X-axis by amino acid start position) for all samples of a given *APOE* genotype in the plasma (**A**) and CSF (**B**), with error bars indicating the SEM. **C,D** Each point represents the average of all peptide values for a given participant in the plasma (**C**) and CSF (**D**), with error bars indicating the SEM.

**Supplementary Figure 3.**
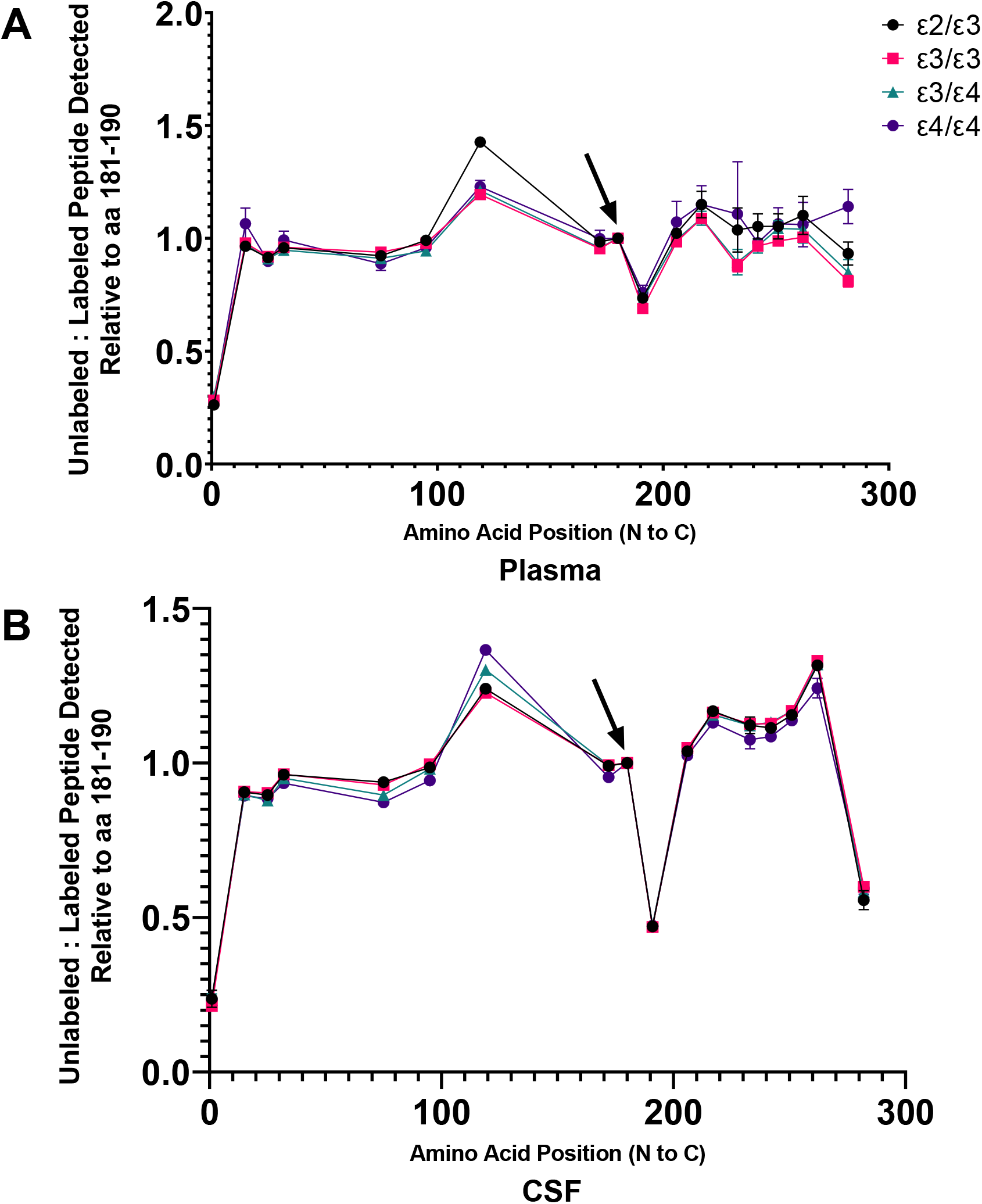
Comparison of the relative amounts of apoE across the unmodified backbone reveal no statistically significant differences at any peptides between *APOE* genotypes in the plasma or in the CSF. Each point represents the mean value for the corresponding peptide relative to the reference peptide, LGPLVEQGR (denoted with a black arrow), in the plasma (**A**) and CSF (**B**), with error bars indicating the SEM, and each color representing a different genotype.

**Supplementary Figure 4.**
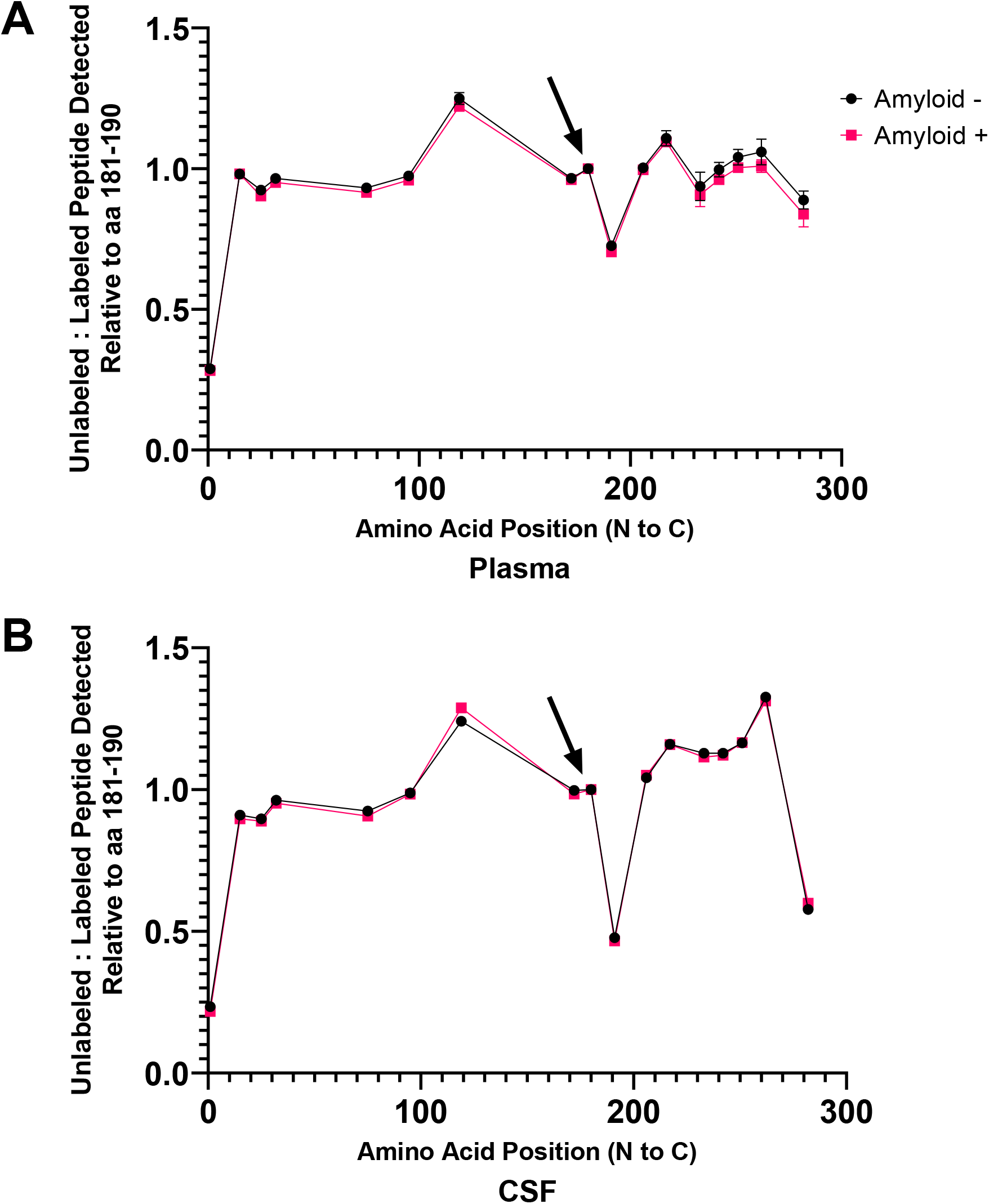
Comparison of the relative amounts of apoE across the unmodified backbone reveal no statistically significant differences across amyloid status in the plasma or in the CSF. Each point represents the mean value for the corresponding peptide relative to the reference peptide, LGPLVEQGR (denoted with a black arrow), in the plasma (**A**) and CSF (**B**), with error bars indicating the SEM.

**Supplementary Figure 5.**
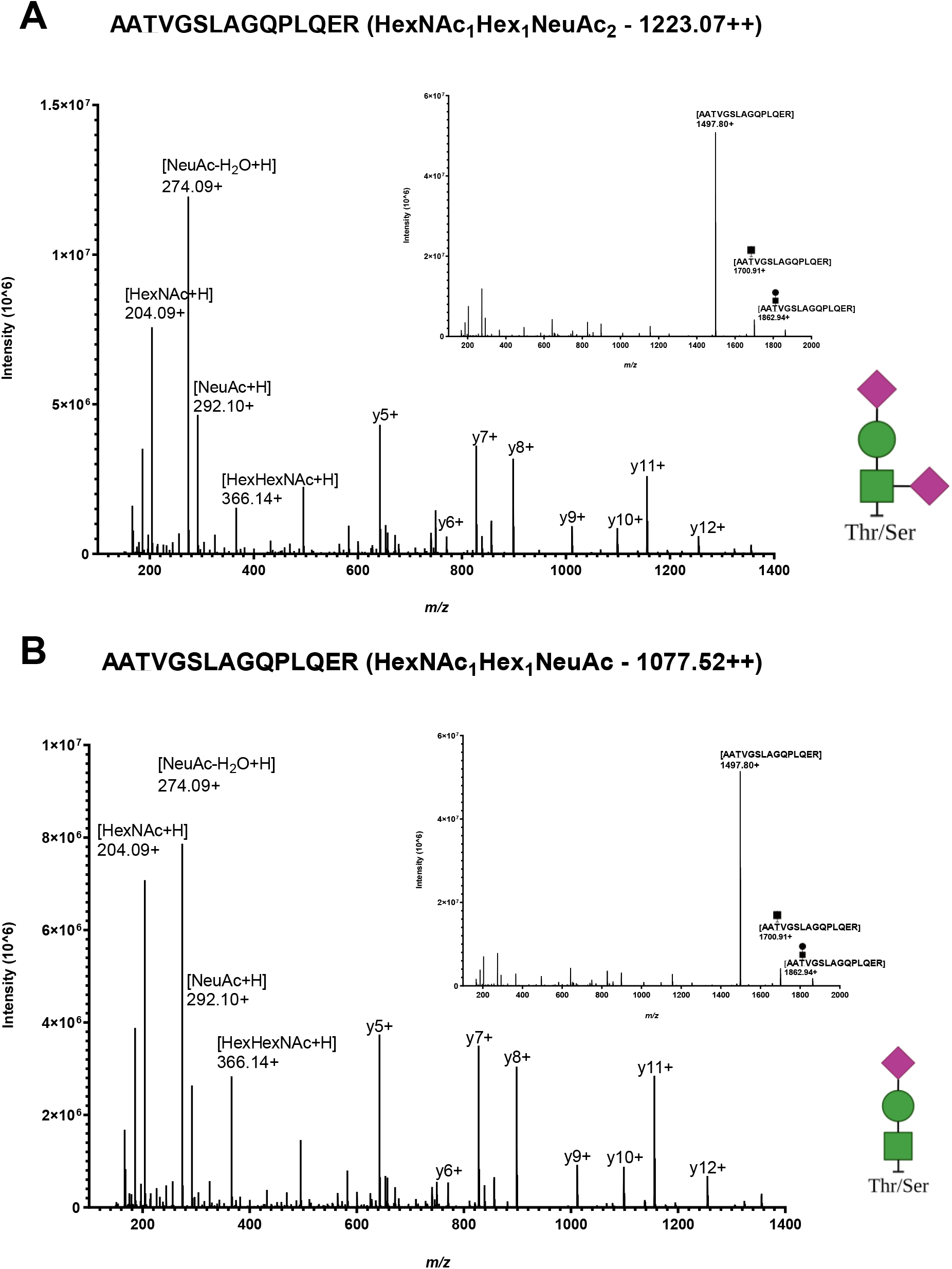
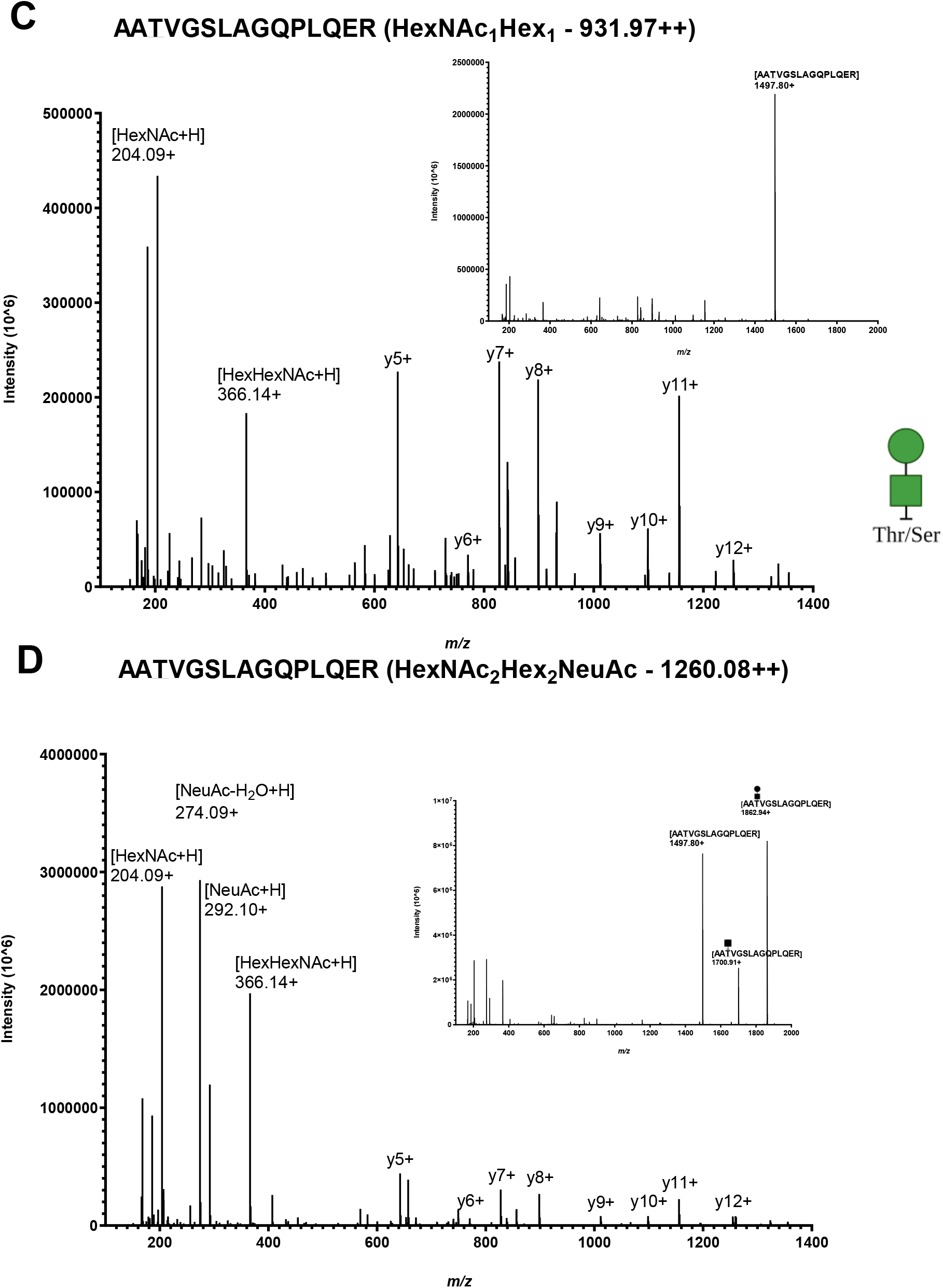
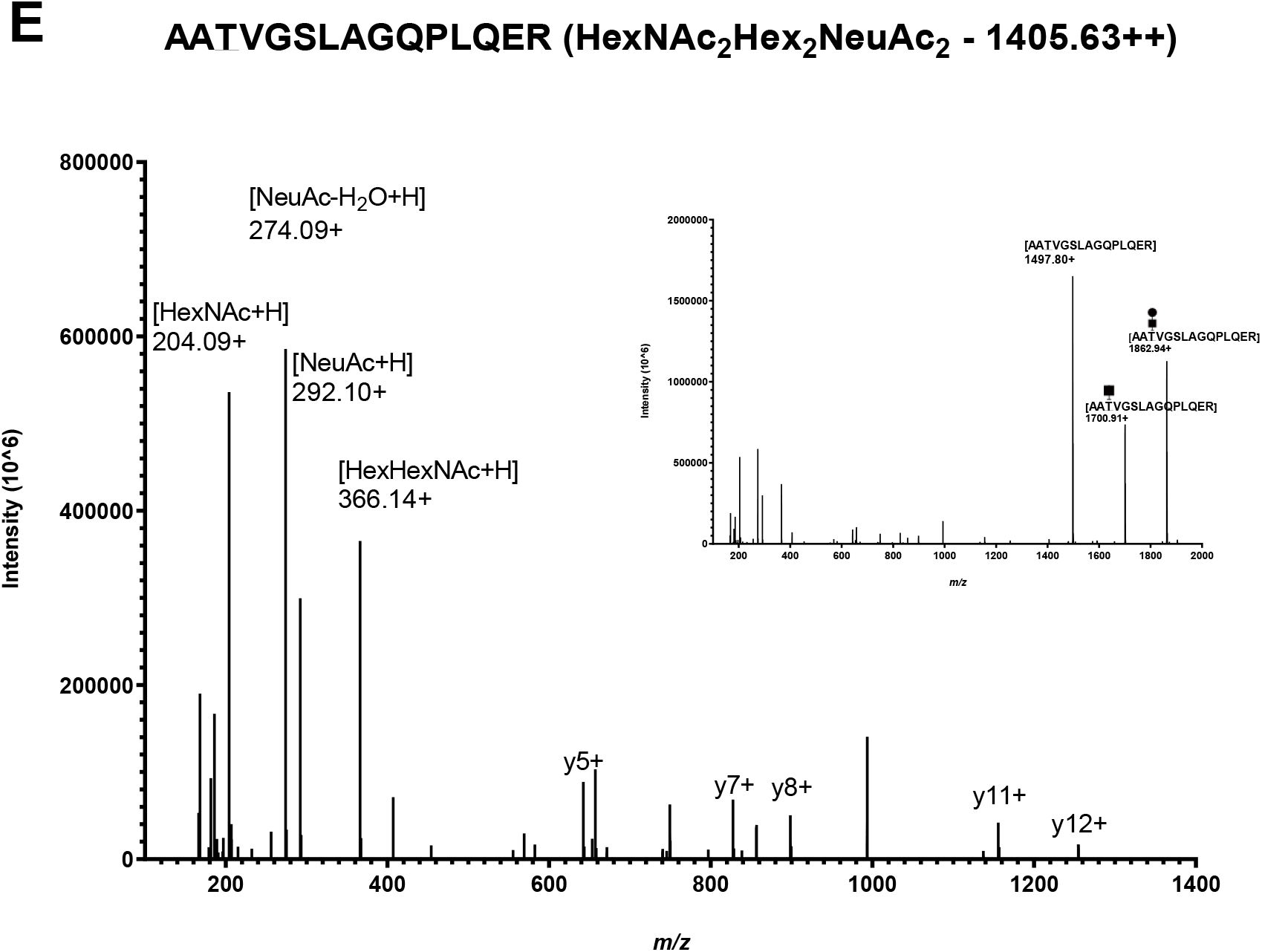
Tandem MS spectra of hinge region core-1 type apoE tryptic glycopeptide AATVGSLAGQPLQER (amino acid residues 192-206) derived from human CSF. **A** Disialylated AATVGSLAGQPLQER. **B** Monosialylated AATVGSLAGQPLQER. **C** Unsialylated AATVGSLAGQPLQER. **D, E** Unsolved AATVGSLAGQPLQER structures.

**Supplementary Figure 6.**
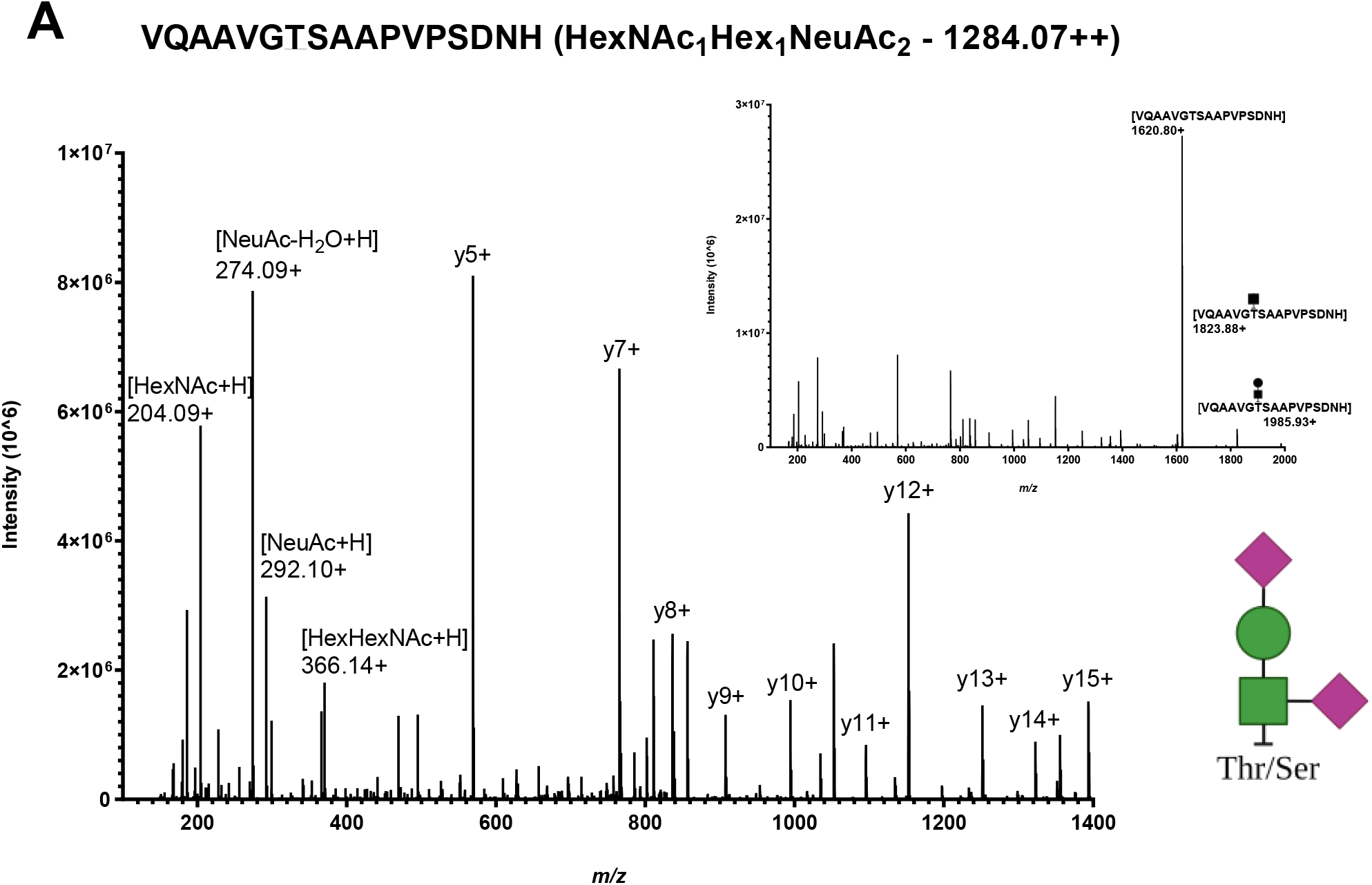
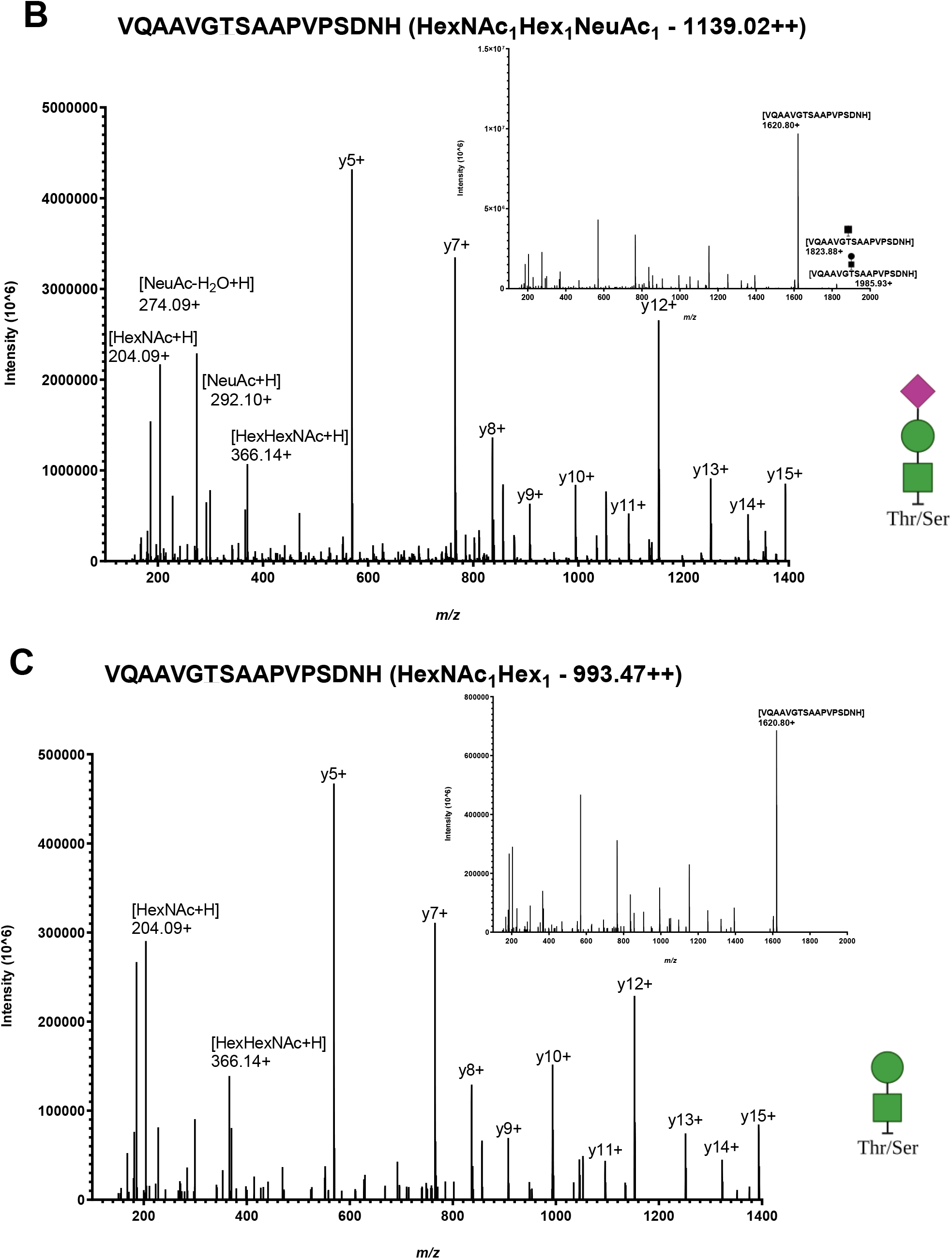
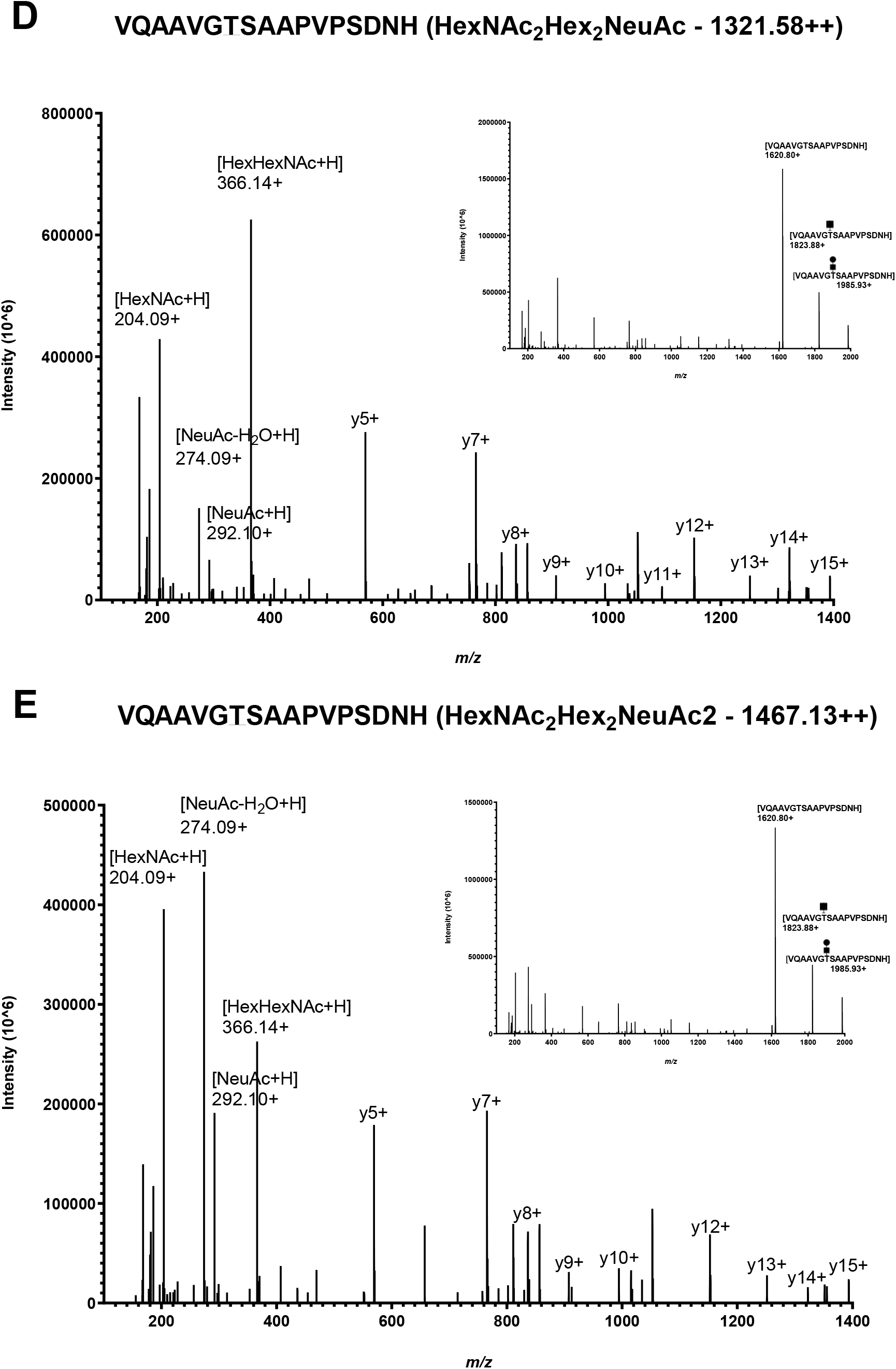
Tandem MS spectra of C-terminal core-1 type apoE tryptic glycopeptide VQAAVGTSAAPVPSDNH (amino acid residues 283-299) derived from human CSF. **A** Disialylated VQAAVGTSAAPVPSDNH. **B** Monosialylated VQAAVGTSAAPVPSDNH. **C** Unsialylated VQAAVGTSAAPVPSDNH. **D,E** Unsolved VQAAVGTSAAPVPSDNH structures.

**Supplementary Figure 7.**
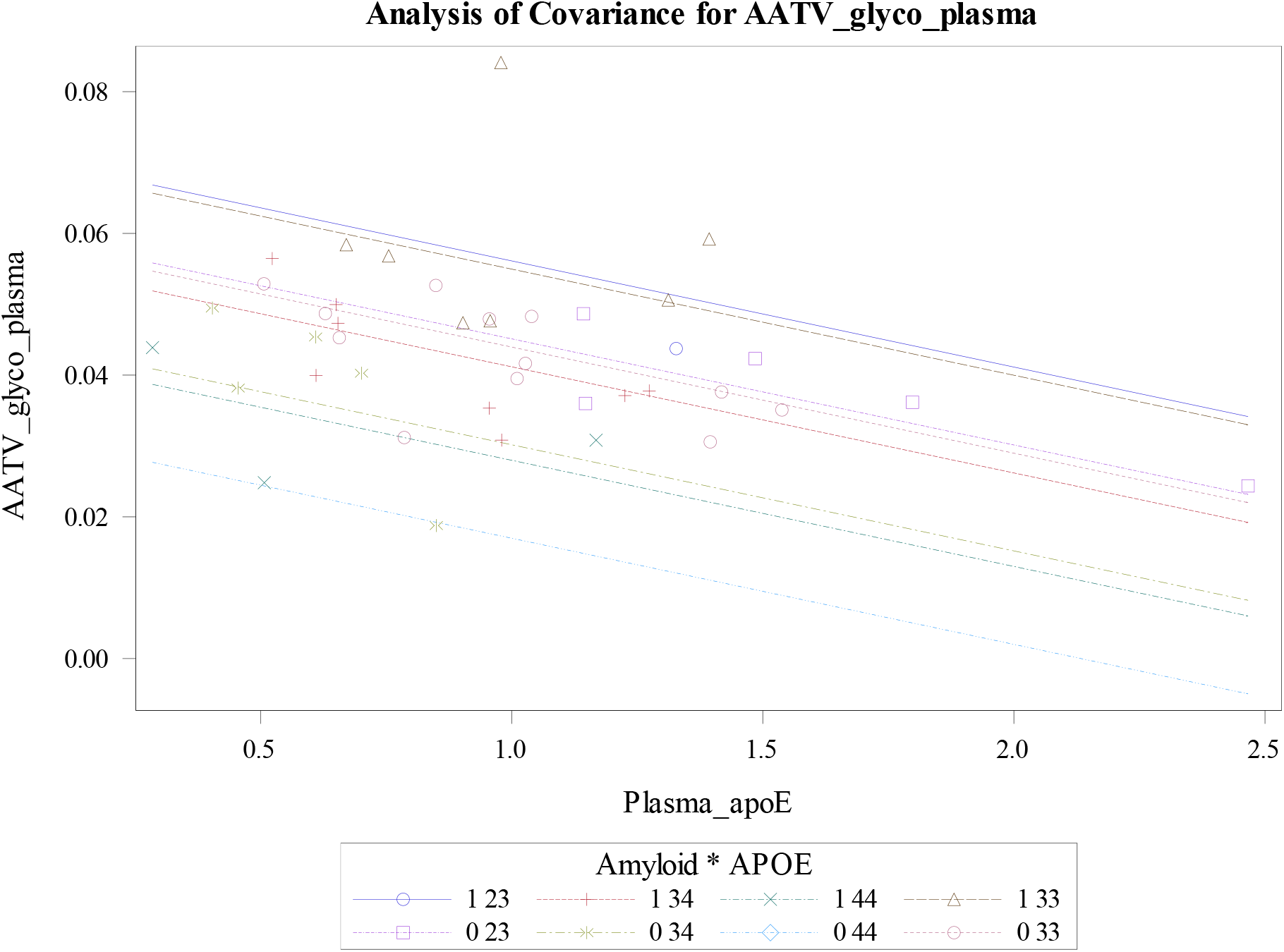
ANCOVA plot of the covariance of plasma total O-glycosylation site occupancy of the hinge region peptide with plasma total apoE amount, *APOE* genotype, and amyloid status. Amyloid status is listed first in the legend (“1” denotes amyloid positive and “0” denotes amyloid negative), followed by *APOE* genotype (“23,” “33,” “34,” “44”).

**Supplementary Table 4.** ANCOVA of plasma O-glycosylation site occupancy for the three predominant hinge region glycopeptides reveals that the unsialylated core 1 and the monosialylated core 1 species demonstrate the strongest correlation with plasma total apoE amount, *APOE* ε4 carrier status and amyloid status. **A** ANCOVA with unsialylated AATVGSLAGQPLQER as the outcome variable. **B** ANCOVA with monosialylated AATVGSLAGQPLQER as the outcome variable. **C** ANCOVA with disialylated AATVGSLAGQPLQER as the outcome variable.

| Parameter | Estimate | SE | *p* = |
| --- | --- | --- | --- |
| Intercept | 0.009 | 0.004 | 0.035 |
| Plasma_apoE | -0.003 | 0.0008 | 0.0003 |
| *APOE* ε4 + | -0.003 | 0.0007 | 0.0006 |
| *APOE* ε4 - | 0 | . | . |
| Amyloid + | 0.002 | 0.0006 | 0.007 |
| Amyloid - | 0 | . | . |
| SEX-M | -0.0006 | 0.0006 | 0.33 |
| SEX-F | 0 | . | . |
| AGE | 0.00001 | 0.00005 | 0.77 |

**B**
| Parameter | Estimate | SE | *p* = |
| --- | --- | --- | --- |
| Intercept | 0.020 | 0.011 | 0.089 |
| Plasma_apoE | -0.007 | 0.002 | 0.002 |
| *APOE* ε4 + | -0.007 | 0.002 | 0.0007 |
| *APOE* ε4 - | 0 | . | . |
| Amyloid + | 0.005 | 0.001 | 0.011 |
| Amyloid - | 0 | . | . |
| SEX-M | -0.0002 | 0.002 | 0.90 |
| SEX-F | 0 | . | . |
| AGE | 0.0001 | 0.0001 | 0.41 |

**C**
| Parameter | Estimate | SE | *p* = |
| --- | --- | --- | --- |
| Intercept | 0.007 | 0.006 | 0.21 |
| Plasma_apoE | -.003 | 0.001 | 0.013 |
| *APOE* ε4 + | -.003 | 0.001 | 0.012 |
| *APOE* ε4 - | 0 | . | . |
| Amyloid + | 0.001 | 0.001 | 0.19 |
| Amyloid - | 00 | . | . |
| SEX-M | 0.0007 | 0.0008 | 0.36 |
| SEX-F | 0 | . | . |
| AGE | 0.0001 | 0.00007 | 0.18 |

**Supplementary Table 5.**
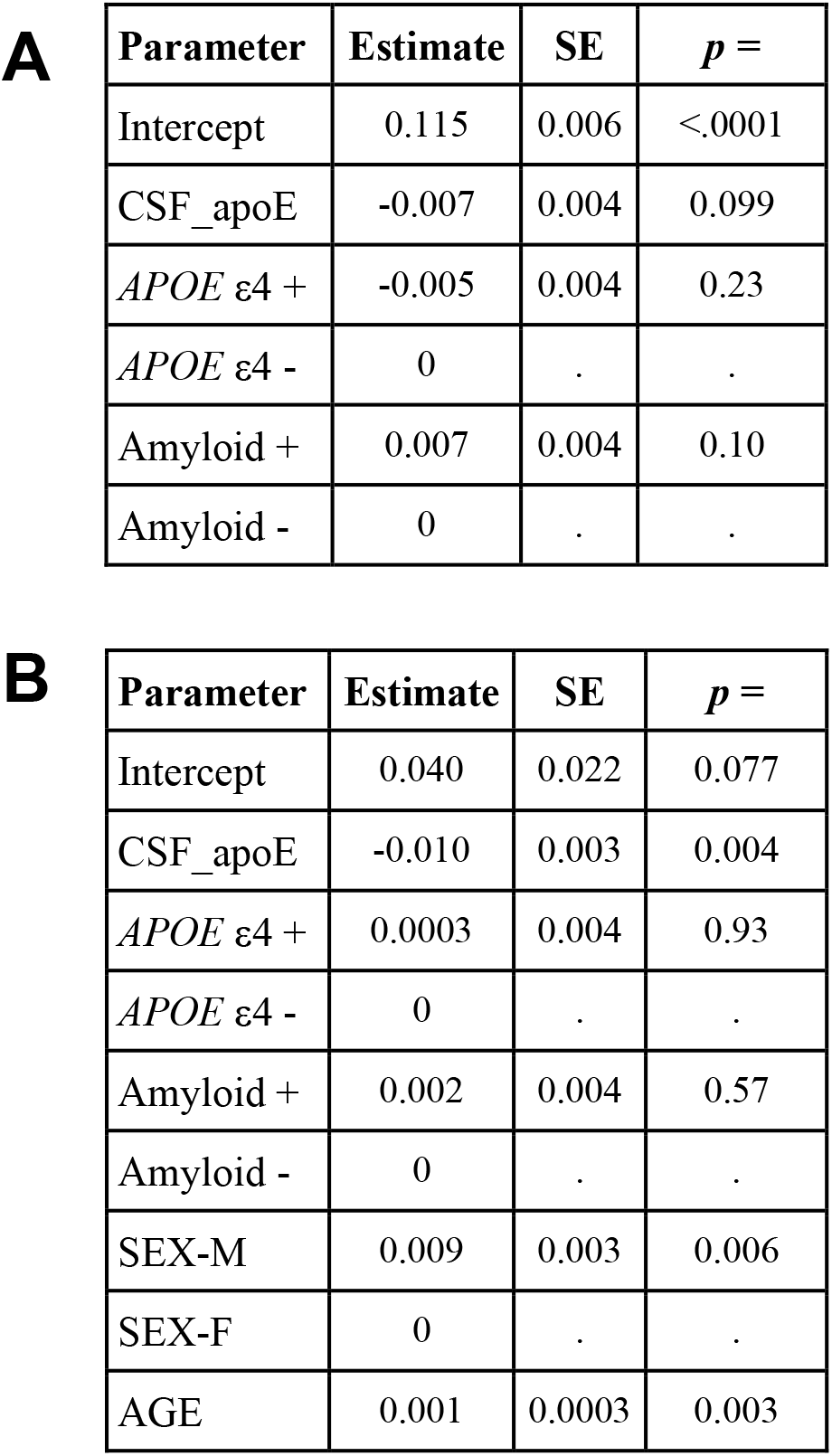
ANCOVA of CSF total O-glycosylation site occupancy of the hinge region peptide reveals a correlation with both sex (p = 0.006) and age (p = 0.003) but no correlation with *APOE* ε4 carrier status, or amyloid status. **A** Total site occupancy as the outcome variable with CSF total apoE as a continuous covariate and *APOE* ε4 carrier status and amyloid status as categorical covariates. **B** Total site occupancy as the outcome variable with CSF total apoE as a continuous covariate and *APOE* ε4 carrier status, amyloid status, sex and age as covariates.

**A**
| Parameter | Estimate | SE | *p* = |
| --- | --- | --- | --- |
| Intercept | 0.115 | 0.006 | <.0001 |
| CSF_apoE | -0.007 | 0.004 | 0.099 |
| *APOE* ε4 + | -0.005 | 0.004 | 0.23 |
| *APOE* ε4 - | 0 | . | . |
| Amyloid + | 0.007 | 0.004 | 0.10 |
| Amyloid - | 0 | . | . |

**Supplementary Table 5 ANCOVA of CSF total O-glycosylation site occupancy of the hinge region peptide reveals a correlation with both sex (*p* = 0.006) and age (*p* = 0.003) but no correlation with *APOE* ε4 carrier status, or amyloid status.**
| Parameter | Estimate | SE | *p* = |
| --- | --- | --- | --- |
| Intercept | 0.040 | 0.022 | 0.077 |
| CSF_apoE | -0.010 | 0.003 | 0.004 |
| *APOE* ε4 + | 0.0003 | 0.004 | 0.93 |
| *APOE* ε4 - | 0 | . | . |
| Amyloid + | 0.002 | 0.004 | 0.57 |
| Amyloid - | 0 | . | . |
| SEX-M | 0.009 | 0.003 | 0.006 |
| SEX-F | 0 | . | . |
| AGE | 0.001 | 0.0003 | 0.003 |

**Supplementary Table 6.**
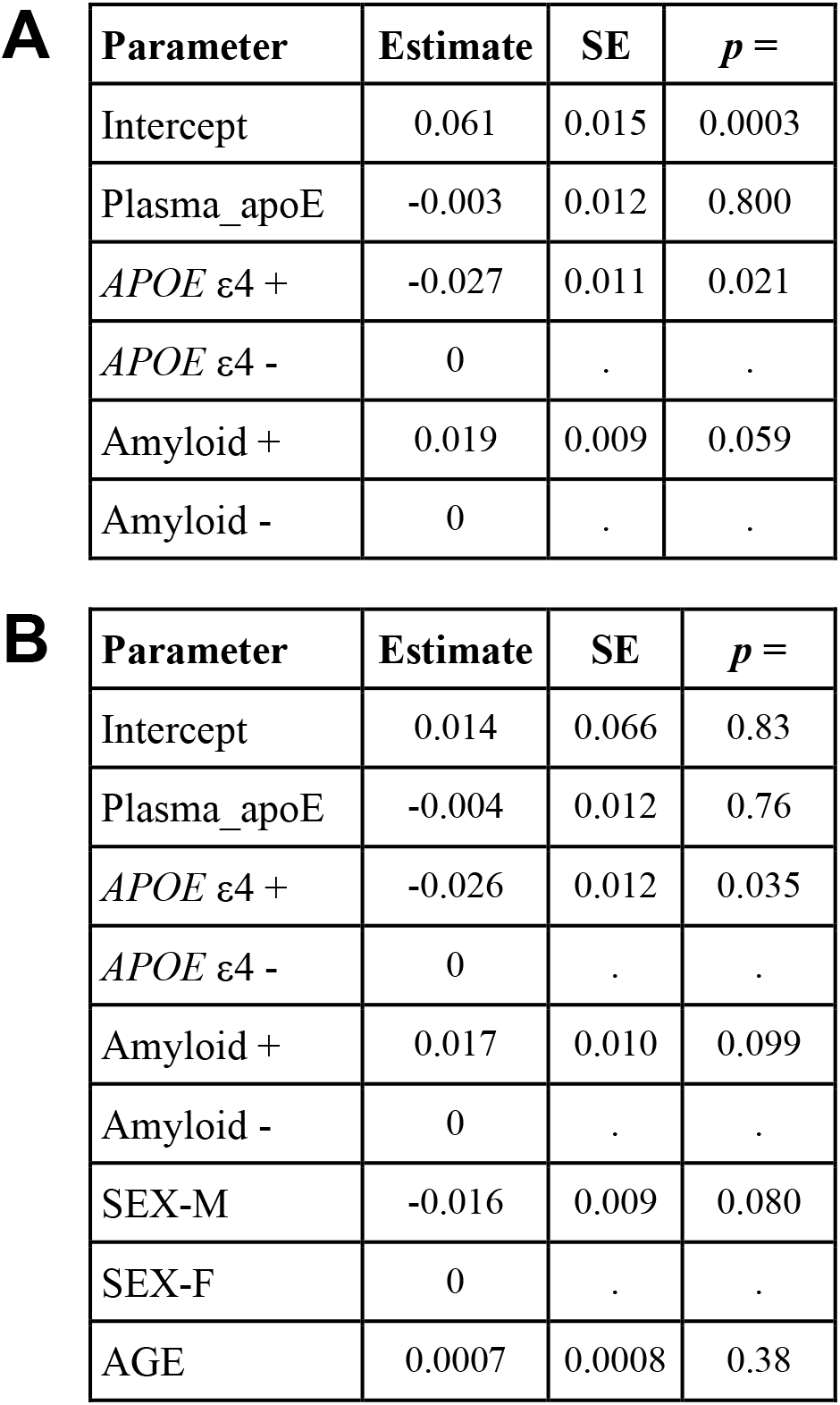
ANCOVA of plasma total O-glycosylation site occupancy of the C-terminal region peptide reveals no correlation with plasma total apoE, *APOE* ε4 carrier status or amyloid status. **A** Total site occupancy as the outcome variable with plasma total apoE as a continuous covariate and *APOE* ε4 carrier status and amyloid status as categorical covariates. **B** Total site occupancy as the outcome variable with plasma total apoE as a continuous covariate and *APOE* ε4 carrier status, amyloid status, sex and age as covariates.

**A**
| Parameter | Estimate | SE | *p* = |
| --- | --- | --- | --- |
| Intercept | 0.061 | 0.015 | 0.0003 |
| Plasma_apoE | -0.003 | 0.012 | 0.800 |
| *APOE* ε4 + | -0.027 | 0.011 | 0.021 |
| *APOE* ε4 - | 0 | . | . |
| Amyloid + | 0.019 | 0.009 | 0.059 |
| Amyloid - | 0 | . | . |

**Supplementary Table 6 ANCOVA of plasma total O-glycosylation site occupancy of the C-terminal region peptide reveals no correlation with plasma total apoE, *APOE* ε4 carrier status or amyloid status.**
| Parameter | Estimate | SE | *p* = |
| --- | --- | --- | --- |
| Intercept | 0.014 | 0.066 | 0.83 |
| Plasma_apoE | -0.004 | 0.012 | 0.76 |
| *APOE* ε4 + | -0.026 | 0.012 | 0.035 |
| *APOE* ε4 - | 0 | . | . |
| Amyloid + | 0.017 | 0.010 | 0.099 |
| Amyloid - | 0 | . | . |
| SEX-M | -0.016 | 0.009 | 0.080 |
| SEX-F | 0 | . | . |
| AGE | 0.0007 | 0.0008 | 0.38 |

**Supplementary Table 7.**
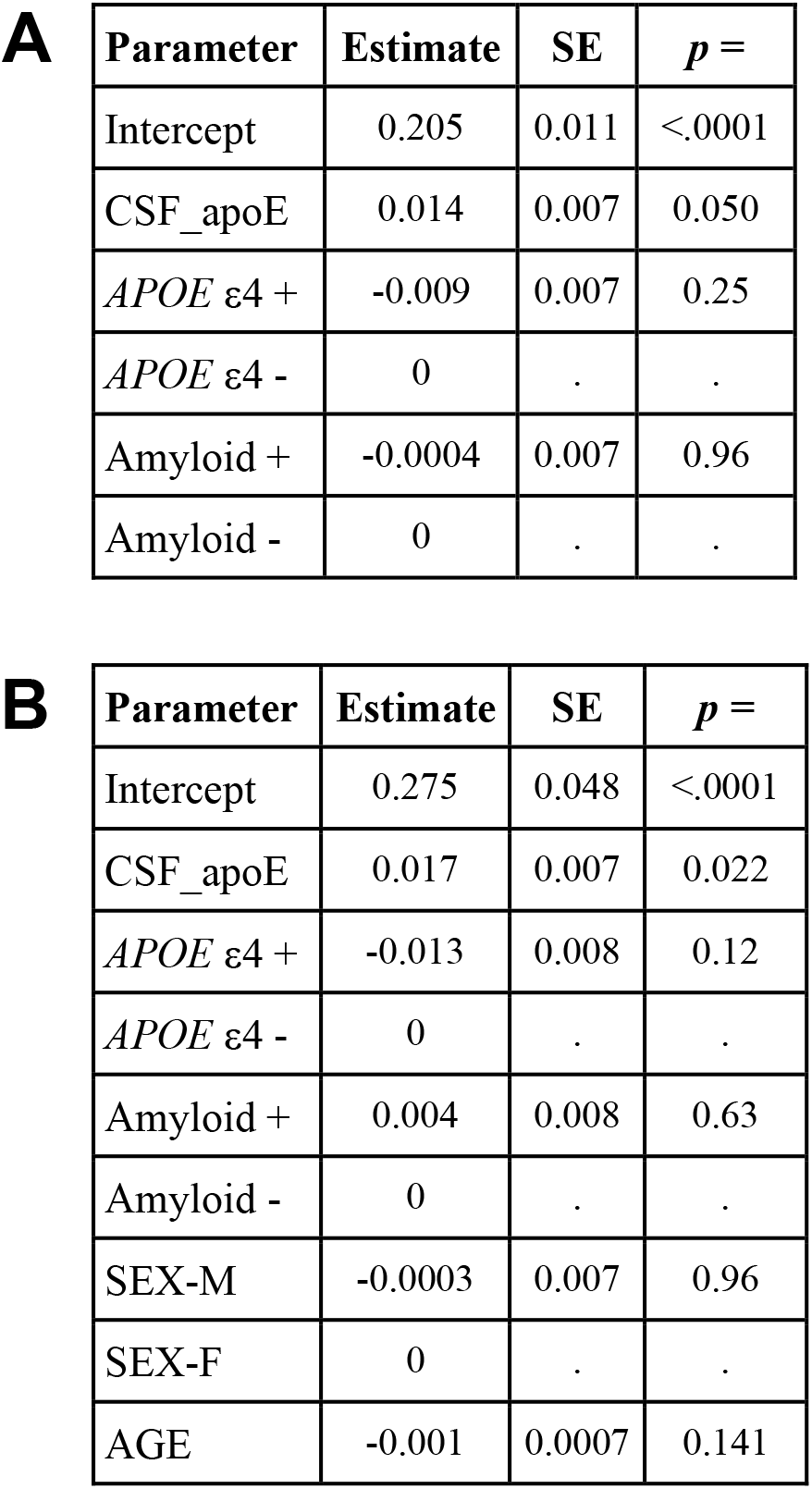
ANCOVA of CSF total O-glycosylation site occupancy of the C-terminal region peptide reveals no correlation with CSF total apoE, *APOE* ε4 carrier status or amyloid status. **A** Total site occupancy as the outcome variable with CSF total apoE as a continuous covariate and *APOE* ε4 carrier status and amyloid status as categorical covariates. **B** Total site occupancy as the outcome variable with CSF total apoE as a continuous covariate and *APOE* ε4 carrier status, amyloid status, sex and age as covariates.

**A**
| Parameter | Estimate | SE | *p* = |
| --- | --- | --- | --- |
| Intercept | 0.205 | 0.011 | <.0001 |
| CSF_apoE | 0.014 | 0.007 | 0.050 |
| *APOE* ε4 + | -0.009 | 0.007 | 0.25 |
| *APOE* ε4 - | 0 | . | . |
| Amyloid + | -0.0004 | 0.007 | 0.96 |
| Amyloid - | 0 | . | . |

**Supplementary Table 7 ANCOVA of CSF total O-glycosylation site occupancy of the C-terminal region peptide reveals no correlation with CSF total apoE, *APOE* ε4 carrier status or amyloid status.**
| Parameter | Estimate | SE | *p* = |
| --- | --- | --- | --- |
| Intercept | 0.275 | 0.048 | <.0001 |
| CSF_apoE | 0.017 | 0.007 | 0.022 |
| *APOE* ε4 + | -0.013 | 0.008 | 0.12 |
| *APOE* ε4 - | 0 | . | . |
| Amyloid + | 0.004 | 0.008 | 0.63 |
| Amyloid - | 0 | . | . |
| SEX-M | -0.0003 | 0.007 | 0.96 |
| SEX-F | 0 | . | . |
| AGE | -0.001 | 0.0007 | 0.141 |

**Supplementary Figure 8.**
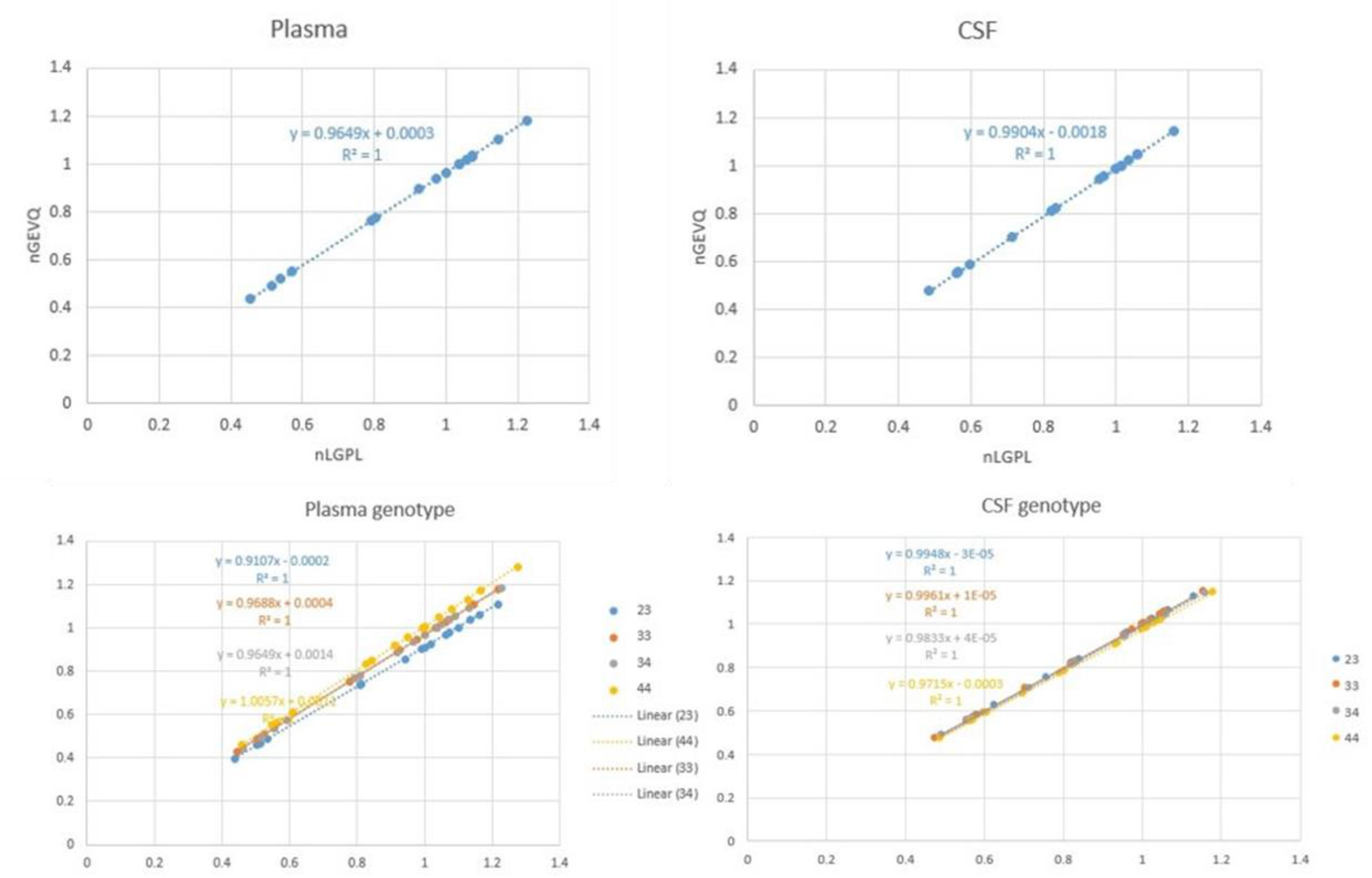
Correlation plots between two peptides, LGPLVEQGR and GEVQAMLGQSTEELR, used as reference peptides for the relative quantification of the apoE peptide backbone.

